# Polyelectrolyte domains in intrinsically disordered regions differentially regulate intracellular biomolecular condensation

**DOI:** 10.1101/2025.09.09.675273

**Authors:** Yoon Jeong Choi, Yujin Lee, Soojin Cho, Yaqi Pan, Yemuk Kang, Changi Pack, Youngtae Jeong, Kiwon Song

## Abstract

Intracellular condensation is a hallmark of many proteins containing intrinsically disordered regions (IDRs). Decoding the regulatory mechanisms of condensation encoded within the amino acid sequences of condensate-forming proteins is essential for understanding both how intracellular condensates assemble and how they contribute to various disease pathologies. Here, we investigated the budding yeast P-body proteins Nst1 and Edc3, which exhibit pronounced intracellular self-condensation, and identified charged amino acid clusters in the low complexity regions (LCRs) that modulate their phase behavior. Guided by the stickers-and-spacers framework, we hypothesized that clusters of charged residues within IDRs function in promoting intracellular condensation process. Using a sliding-window approach, we designated and dissected the polyelectrolyte (PE) domains of Nst1 and Edc3 and established their essential contributions to condensate coalescence. In Nst1, the polyanionic (PA) subdomain enhances intermolecular attraction and promotes coalescence, whereas the polyampholyte (PAM) subdomain imparts rigidity to the condensate as a distinct modulatory element and functions as a platform in condensation process of other P-body components. PA of Edc3 also regulates coalescence. Extending these findings to the neurodegenerative disease-associated protein TDP-43, we show that its PA regions are critical for coalescence, consistent with the presence of negatively charged amino acid clusters as potent molecular stickers functioning in condensate coalescence in cells. These results highlight charged-residue clusters as important sequence determinants of protein phase behavior and suggest that they represent promising novel regulatory targets for modulating TDP-43 condensation in cells.

## INTRODUCTION

Biomolecular condensation observed in cells not only deepens our understanding of molecular dynamics in response to various physiological conditions, but also shed light on the mechanisms underlying protein amyloid formation (Alberti and Hyman, 2021; Alshareedah et al., 2024; Banani et al., 2017; Niu et al., 2023). These condensations occur within the crowded and complex cellular environment, exhibiting diverse physical properties depending on the degree of condensation.

Among these behaviors, liquid-like demixing has been extensively studied as one prominent mechanism of condensate formation, in which dynamic, droplet-like assemblies are distinguished from more rigid, amyloid-like protein aggregates (Ranganathan et al., 2022). Neurodegeneration-associated proteins usually act as scaffolds that drive the formation of liquid-like condensates with self-condensation ability and high multivalency. Recent studies have increasingly reported the co-occurrence of amyloid fibrils and liquid-like condensates of pathogenic proteins, such as TDP-43, both in the brain tissues of patients with neurodegenerative diseases and in cellular models of these diseases (Ahmad et al., 2022; Alshareedah *et al*., 2024; Gruijs da Silva et al., 2022; Sneideriene et al., 2025). This observation strongly suggests that the aggregation of pathogenic proteins does not proceed through direct solid aggregation, but rather through prior enrichment into liquid-like condensates, followed by a structural conversion into amyloid assemblies. In other words, dysregulation of these liquid-like condensates is expected to promote the formation of protein amyloids implicated in neurodegenerative disorders (Hurtle et al., 2023; Nam and Gwon, 2023; Visser et al., 2024). Therefore, elucidating the underlying mechanisms of liquid-like condensation is critical both for understanding the molecular dynamics underlying physiological condensate formation, and for discovering therapeutic strategies to regulate pathological condensate formation (Alberti and Hyman, 2021; Choi et al., 2024; Gao et al., 2023; Lu et al., 2024; Mehta and Zhang, 2022).

Stickers-and-spacers model is a leading framework of a molecular mechanism for biomolecular liquid-like condensation (Banjade and Rosen, 2014; Choi et al., 2020; Sood and Zhang, 2024). This model defines “stickers” as molecular elements that mediate attractive interactions, and “spacers” as flexible linkers that separate stickers and modulate condensate properties (Choi *et al*., 2020). Amino acids in intrinsically disordered regions (IDRs), capable of functioning as stickers, were precisely identified through *in vitro* studies. The cation-*π* interaction between tyrosine and arginine, as well as the interaction between lysine and RNA, were revealed to act as stickers (Ukmar-Godec et al., 2019; Wang et al., 2018b). The sequential pattern of these components significantly influences the dynamic characteristics of the condensate (Alshareedah *et al*., 2024; Alshareedah et al., 2021; Wang *et al*., 2018b). Notably, when sticker amino acids are clustered within the sequence, rather than dispersed, the resultant condensate tends towards solidification (Alshareedah *et al*., 2024; Alshareedah *et al*., 2021; Wang *et al*., 2018b). Thus, the stickers-and-spacers model offers a solid basis for elucidating the condensation characteristics of molecules *in vitro*, offering valuable insights into the behavior of biomolecular condensates.

Despite the explanatory power of the stickers-and-spacers model, the molecular grammar underlying intracellular biomolecular condensation remains enigmatic. First, the morphology and physical properties of condensates depend on the characteristics of condensate-forming proteins and the intracellular environment, and these properties can change over time as condensation progresses. Thus, the stickers that function may vary depending on the phase of condensates. Second, although the molecular grammar of condensation has been extensively characterized for prion-like low-complexity domains (PLCDs) - such as poly Q/N tracts in yeast amyloidogenic proteins and the RGG motifs in disease-related proteins like FUS and TDP-43 (Patel et al., 2015; Sanders et al., 2020; Wang et al., 2018), other intrinsically disordered regions (IDRs) with distinct combinations of sticker amino acids remain largely unexplored. Critically, many IDRs lack canonical PLCD features but still drive phase separation, suggesting the existence of alternative molecular grammars.

Recent *in vitro* and simulation studies have reported that the charge distribution within amino acid sequences strongly affects the viscosity, surface tension, and dynamic properties of condensates (Biswas and Potoyan, 2024; Linsenmeier et al., 2023; Sundaravadivelu Devarajan et al., 2024). These reports strongly indicate the need for research on the regulation of cellular condensation process by novel low complexity regions (LCRs) composed of sticker amino acids such as charged amino acid clusters, in addition to the previously studied PLCDs.

In this study, we aimed to elucidate the role of the polyelectrolyte region in the cellular condensation of biomolecules via Nst1. Both Nst1 and Edc3, established P-body components in budding yeast, possess intrinsic self-condensation capabilities. Upon Nst1 and Edc3 overexpression, cells initially exhibit numerous small condensates of Nst1 and Edc3, which progressively fuse into larger structures as protein levels accumulate over time, thereby reducing the overall number of observed condensate (Fig. 1A). Utilizing this model, we examined how the polyelectrolyte region affects both the molecular dynamics of NST1 condensates and the condensate formation of P-body components mediated by NST1. Furthermore, we investigated whether the function of the polyelectrolyte region in Nst1 condensation could be extrapolated to TDP-43, a stress granule constituent that is closely linked to the pathology of neurodegenerative diseases.

**Figure 1.**
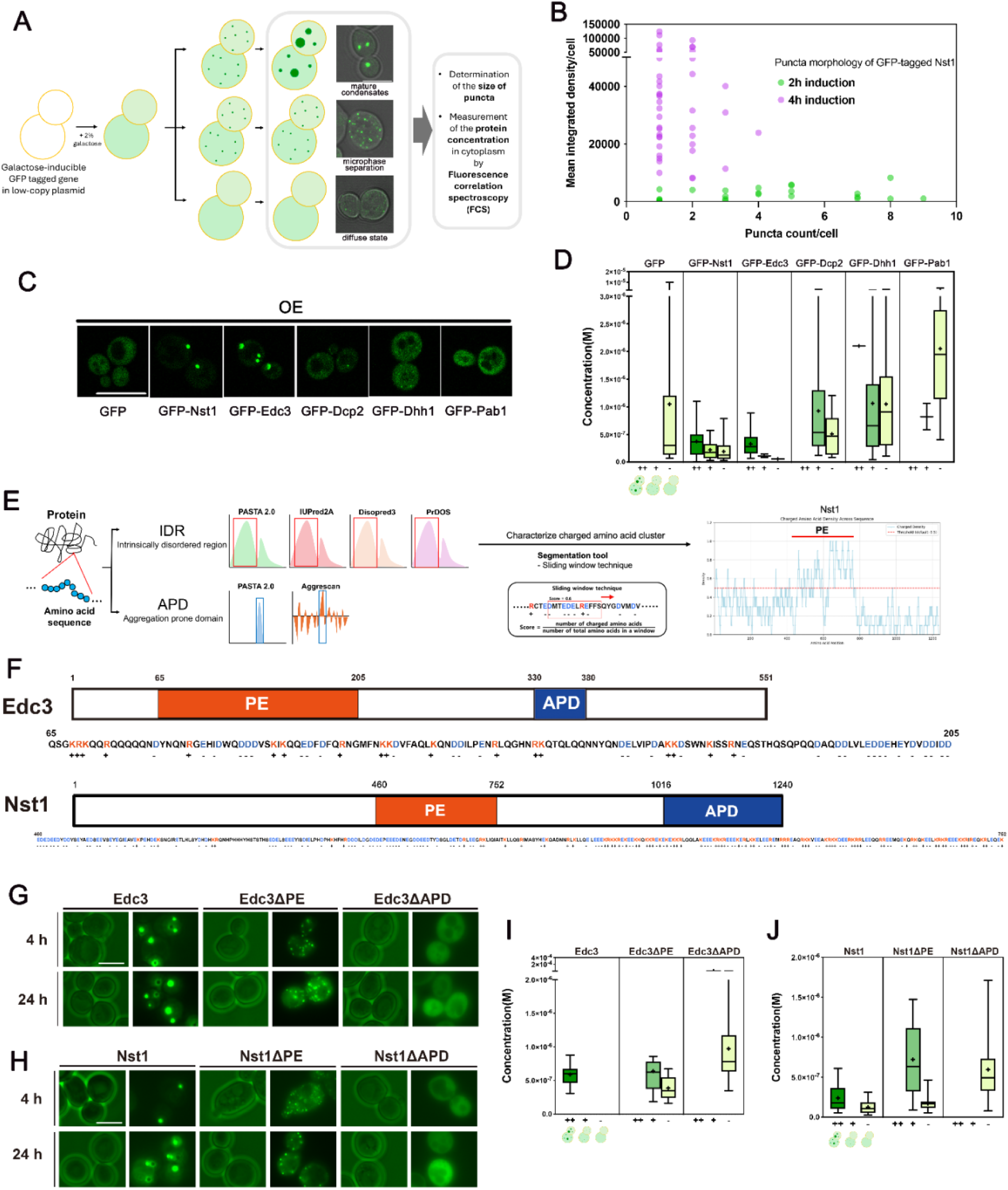
Overexpression of Nst1 and Edc3 leads to mature condensate formation, whereas overexpression of their polyelectrolyte (PE) domain deletion mutants within the IDR maintains a stabilized microphase-separated state. (A) Schematic representation of budding yeast overexpression system to observe intracellular condensates. (B) Diagram of puncta generation pattern of cells overexpressing GFP tagged Nst1 for 2 h (green) and 4 h (purple). *Δnst1* (YSK3490) cells transformed with pMW20-pGAL-GFP-NST1 were induced for 2 hours (green) and 4 hours (purple) under the GAL promoter. Cells were imaged and analyzed; mean integrated density and the number of puncta of each cell were measured and plotted. (C-D) Representative fluorescence microscopy images and quantitative phase separation analysis of cells overexpressing GFP and GFP-tagged Nst1, Edc3, Dcp3, Dhh1, and Pab1. (C) Scale bar: 10 μm. (D) Cytoplasmic protein concentrations were measured via fluorescence correlation spectroscopy (FCS) as described in Materials and Methods. The puncta formation patterns in cells were classified into three categories: diffuse phase, microphase separation, and mature condensates. Box plot analysis of the correlation between the pattern of protein puncta formation and cytoplasmic concentration was presented. (E) Schematic representation of polyelectrolyte (PE) domain segmentation. (F) Sequence diagram of Nst1 and Edc3. The red box indicates the polyelectrolyte (PE) region, while the blue box represents the aggregation-prone domain (APD). Each sequence shown corresponds to the PE region. Positively charged amino acids, lysine (K) and arginine (R),, are highlighted in red and marked with a (+) sign below, whereas negatively charged amino acids, glutamic acid (E) and aspartic acid (D), are highlighted in blue and marked with a (–) sign below. (G) Fluorescence microscopy images showing puncta formation in cells overexpressing Edc3 and each domain deletion mutant after 4 and 24 hours of galactose induction. In *Δedc3* cells (YSK3707), GFP-tagged full-length Edc3 (pMW20-pGAL-GFP-EDC3), Edc3ΔPE (pMW20-pGAL-GFP-EDC3ΔPE), and Edc3ΔAPD (pMW20-pGAL-GFP-EDC3ΔAPD) were overexpressed. Representative images were selected from an average of more than 100 cells per strain at each time point. Scale bar: 5 µm. (H) Fluorescence microscopy images showing puncta formation in cells expressing Nst1 and each Nst1 domain deletion mutant after 4 and 24 hours of induction as in (E). GFP-tagged full-length Nst1 (pMW20-pGAL-GFP-NST1), Nst1ΔPE (pMW20-pGAL-GFP-NST1ΔPE), and Nst1ΔAPD (pMW20-pGAL-GFP-NST1ΔAPD) were overexpressed in *Δnst1* (YSK3490) cells. Representative images were selected from an average of more than 100 cells per strain at each time point. Scale bar: 5 µm. (I) Box plot analysis of the correlation between the puncta formation pattern and cytoplasmic concentration of the Edc3 and Edc3 derivatives. The puncta formation patterns in cells were classified into three categories: diffuse phase, microphase separation, and mature condensates. Cytoplasmic concentrations were measured in cells expressing each Edc3 domain deletion mutant after 4 hours of induction by fluorescence correlation spectroscopy (FCS) as described in Materials and Methods. The measured cytoplasmic concentrations of cells were analyzed with respect to puncta formation patterns. (J) Box plot analysis of the correlation between the puncta formation pattern and cytoplasmic concentration of the Nst1 and Nst1 derivatives. The cytoplasmic protein concentration measurement and puncta formation analysis were performed same as (F).

## RESULTS

### Overexpression of Nst1 and Edc3 leads to mature condensate formation, whereas overexpression of their polyelectrolyte (PE) domain deletion mutants within the IDR maintains stabilized microphase-separated state

In our previous study, we reported that GFP-tagged Nst1 and Edc3, when overexpressed under the control of the GAL promoter, formed condensates that were larger and denser than those formed by Dcp2 or Dhh1 (Choi et al., 2022). Here, we monitored the formation dynamics of these condensates from the early phase of Nst1 and Edc3 overexpression until their intracellular expression levels reached saturation (Fig. 1A). Numerous small condensates—here referred to as microphase-separated condensates—began to appear simultaneously throughout the cell as early as 1 hour after galactose induction and were observed in nearly all cells by 2 hours. When we analyze the mean integrated density and the number of condensates of each cell at 4 hours post-induction, the number of condensates per cell had decreased, while their internal density had increased markedly, indicative of a transition to a mature condensate state (Fig. 1B). These observations suggest that microphase Nst1 condensates progressively undergo coalescence as the intracellular concentration of Nst1 rises.

Next, we confirmed this distinctive self-condensation potential of Nst1 and Edc3 through analysis of puncta formation patterns in correlation with cytoplasmic protein concentration (Fig. 1C-D). Each protein exhibits a distinct critical concentration for condensation, and a lower critical concentration indicates a higher potential for molecular condensation. By measuring the cytoplasmic concentration of proteins and correlating it with the degree of puncta formation (microphase separation, mature condensates), we can compare condensation patterns between proteins under cellular conditions. Using a galactose promoter, GFP-tagged P-body components such as Nst1, Edc3, Dcp2, and Dhh1, along with the stress granule component Pab1, were overexpressed for 4 hours. Afterward, the cytoplasmic concentration of the overexpressed proteins and the degree of self-condensation were measured using fluorescence correlation spectroscopy (FCS) (Fig.1 A, C-D). We classified the patterns of puncta formation into three categories: diffuse state, microphase separation, and mature condensates (Fig. 1A). GFP control did not exhibit puncta formation at any measured cytoplasmic concentration, whereas Nst1 and Edc3 demonstrated mature condensates at remarkably low concentrations (Fig. 1C-D). Another P-body component, Dcp2 and Dhh1, exhibited microphase separation at higher cytoplasmic concentrations than the concentrations required for mature condensate formation by Nst1 and Edc3. However, mature condensates similar to those observed with Nst1 and Edc3 were not detected for Dcp2 and Dhh1 (Fig. 1C-D). These results confirm that, compared to Dcp2 and Dhh1, Nst1 and Edc3 have a greater intrinsic propensity for self-condensation in cells, which indicates their intrinsic potential for condensation.

Next, we hypothesized that a particular charged-amino acid cluster within intrinsically disordered regions (IDRs) in Nst1 protein sequence may function as a “sticker” domain during condensate formation. In our previous study, we identified that PAM (polyampholyte domain, previously described as PD), a segment of a charged-amino acid cluster in Nst1, functions in the condensation of other P-body components in cells overexpressing Nst1 (Choi *et al*., 2022). However, Nst1 has extended charged amino acid regions beyond PAM. We systematically searched for polyelectrolyte-like clusters of charged amino acids in the IDRs of Nst1 and Edc3 using a sliding window approach, as described in Supplementary Figure 1E (Fig. 1E). Charged amino acids (K, R, H, D, E) were classified, and their proportion within each window was calculated to generate a scoring system. Subsequently, a density plot of charged amino acids was created based on their positions within the sequence (Fig. 1E, S1, S2). Using this method, we segmented polyelectrolyte (PE) amino acid sequence regions spanning residues 65–205 in Edc3 and 460–752 in Nst1 (Fig. 1F, S1, S2).

**Figure 2.**
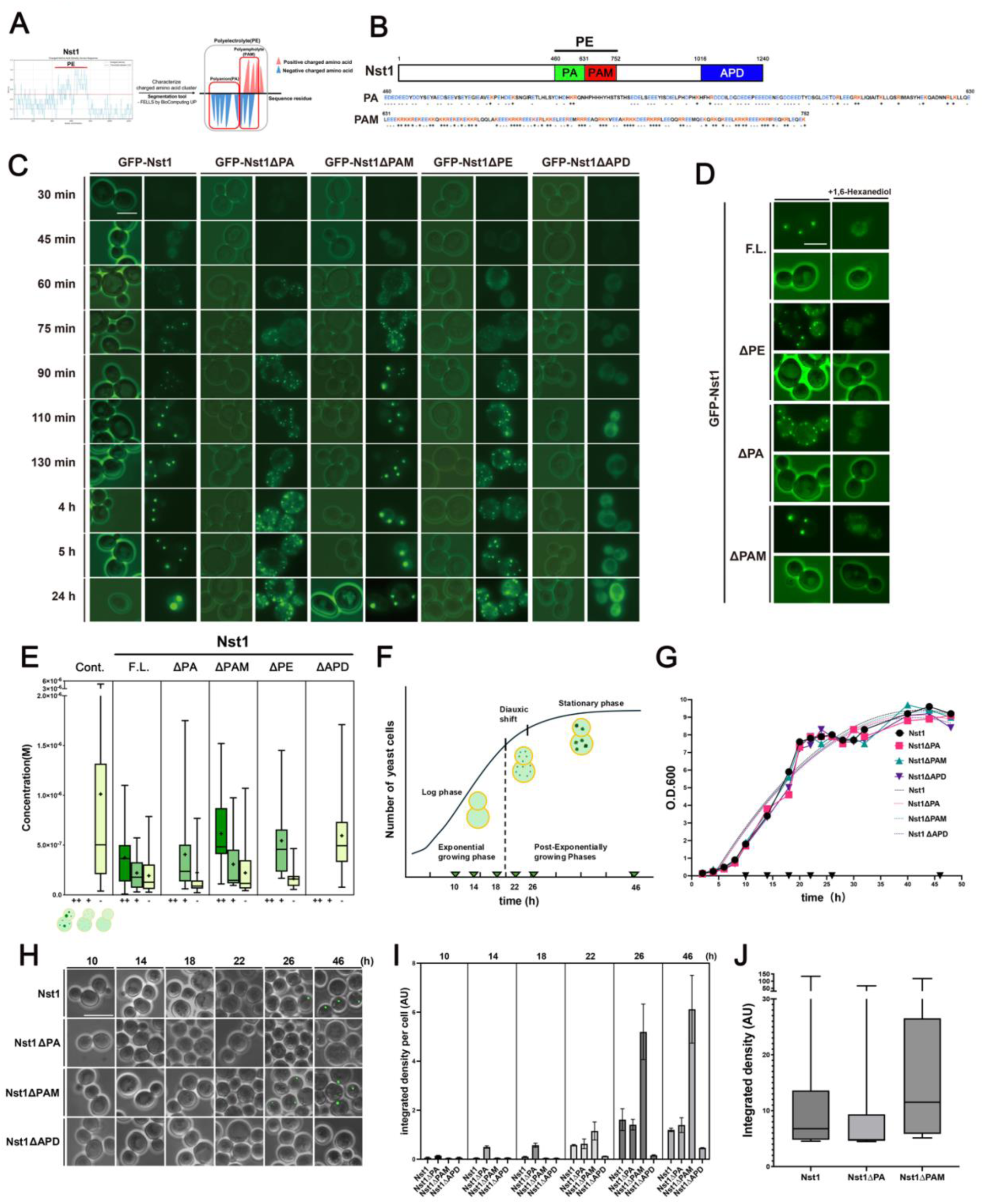
Polyanion (PA) domain of the Polyelectrolyte (PE) region specifically promotes the coalescence of Nst1 condensates, whereas the polyampholyte (PAM) domain in the PE region does not. (A) Schematic representation of Polyanion (PA) and Polyampholyte (PAM) domain segmentation with a sliding window approach. (B) Diagram of Nst1 with its domain architectures and sequences. The black line indicates the PE region, while the green, red, and blue boxes denote the PA, PAM, and APD regions, respectively. The amino acid sequences of the PA and PAM regions are shown below the diagram. Charges were annotated as described in Figure 1D. (C) Fluorescence microscopy images of condensate formation in cells overexpressing each GFP-tagged Nst1 and its domain deletion mutants. Expression of GFP-tagged full-length Nst1 (pMW20-pGAL-GFP-NST1) or NST1 lacking specific domains - Nst1ΔPE (pMW20-pGAL-GFP-NST1ΔPE), Nst1ΔPA (pMW20-pGAL-GFP-NST1ΔPA), Nst1ΔPAM (pMW20-pGAL-GFP-NST1ΔPAM), and Nst1ΔAPD (pMW20-pGAL-GFP-NST1ΔAPD) - was induced in *Δnst1* (YSK3490) cells. Each domain deletion mutant was overexpressed by galactose induction for the indicated time periods. At each time point, fluorescence images were collected from at least 100 cells to ensure representativeness. Scale bar: 5 µm. (D) Fluorescence microscopy images of GFP-tagged Nst1 condensates induced for 4 hours, followed by a 1-hour treatment with 1,6-hexanediol. Scale bar: 5 µm. (E) Box plot analysis of the correlation between the puncta formation pattern and cytoplasmic concentration of cells overexpressing GFP-tagged Nst1, Nst1ΔPA, Nst1ΔPAM, Nst1ΔPE, and Nst1ΔAPD. Cytoplasmic protein concentrations were measured via fluorescence correlation spectroscopy (FCS). The puncta formation patterns in cells were classified into three categories: diffuse phase, microphase separation, and mature condensates. (F-J) Biomolecular condensate formation of endogenously expressed GFP-tagged Nst1, Nst1ΔPA, Nst1ΔPAM and Nst1ΔAPD during chronological aging in yeast. (F) The growth phase diagram of general budding yeast with observation time points designated. The inverted triangles (▾) on the x-axis represent the observed time points (10, 14, 18, 22, 26, and 46 hours). (G) Line chart showing the optical densities (O.D.600) of cell cultures endogenously expressing GFP-tagged Nst1, Nst1ΔPA, Nst1ΔPAM and Nst1ΔAPD under native NST1 promoter. The growth of *Δnst1* (YSK3490) cells integrated with pRS306-pNST1-GFP-NST1 full-length and various NST1 deletion mutants (YSK3700, YSK3701, YSK3702, YSK3714) was monitored over time. The 10 h,14 h and 18 h time points represent the log phase, 22 h the diauxic shift, and 26 h and 46 h the stationary phase. (H) Condensates were segmented using Fiji (ImageJ) and visualized in green. Shown are GFP images merged with corresponding phase-contrast images. Scale bar: 10 μm. (I) Bar plot showing the average integrated GFP fluorescence density per cell for each yeast strain expressing full-length Nst1 and its deletion mutants (Nst1ΔPA, Nst1ΔPAM, Nst1ΔAPD), as shown in (H). For each condition, more than 300 cells were analyzed using Fiji (ImageJ). Results are presented using GraphPad Prism 10. (J) Box plot showing the integrated GFP intensity of condensates in stationary-phase (46 h) yeast cells expressing Nst1, Nst1ΔPA and Nst1ΔPAM. For each condition, more than 300 cells were quantified using Fiji (ImageJ). Data from three independent biological replicates were plotted with GraphPad Prism 10.

Then, we generated deletion mutant clones of Nst1ΔPE and Edc3ΔPE, overexpressed each protein, and observed the cells after 4 hours of induction. Consistent with previous findings, overexpression of Nst1 deleted with the aggregation prone domain (APD), Nst1ΔAPD, did not result in condensate formation (Choi *et al*., 2022) even at cytoplasmic concentrations comparable to those where wild-type Nst1 forms condensates (Fig. 1G-J). Interestingly, cells overexpressing the PE deletion mutants of both Nst1 and Edc3 exhibited microphase separation, even when their cytoplasmic concentrations were comparable with those of wild type Nst1 and Edc3 (Fig. 1G-J). To rule out insufficient expression or time as the cause of microphase separation, we extended the culture to 24 hours in galactose medium. However, even after prolonged induction, both Nst1ΔPE and Edc3ΔPE failed to form mature condensates and continued to exhibit microphase separation (Fig. 1G, H). These results indicate that each polyelectrolyte (PE) is necessary for the coalescence of both Nst1 and Edc3 condensates, thereby facilitating the formation of mature condensates of Nst1 and Edc3.

### Polyanion (PA) domain of the Polyelectrolyte (PE) region specifically promotes the coalescence of Nst1 condensates, whereas the polyampholyte (PAM) domain in the PE region does not

Since the PE region of Nst1 exhibits a unique pattern of charged amino acid arrangement compared to the PE region of Edc3, the PE region of Nst1 was further characterized. Analysis of the Nst1 PE region using FELLS (http://protein.bio.unipd.it/fells/ BioComputing UP) revealed two distinct segments: a polyanion (PA) region spanning residues 460–631, a cluster consisting of remarkably high proportion of negatively charged amino acids, and a polyampholyte (PAM) domain spanning residues 632–752 (Fig. 2 A, B). In our previous study, we reported that the PAM domain (PD in the previous report) is required for promoting the condensation of other P-body components such as Dcp2 and Dhh1, which is induced by Nst1 condensation, but does not affect the self-condensation of Nst1 (Choi *et al*., 2022).

To validate whether the effects of PE on Nst1 condensation were specifically due to the PA domain, we observed the time-dependent pattern of puncta formation of each GFP-tagged Nst1 domain deletion mutant in *Δnst1* cells. When overexpressed for 1 hour using the galactose induction system, the wild-type Nst1 (Nst1 full-length, F.L.), Nst1ΔPA, Nst1ΔPAM, and Nst1ΔPE all exhibited microphase separation (Fig. 2C). After 4 hours of induction, wild-type Nst1 and Nst1ΔPAM exhibited mature condensates with a reduced number of puncta per cell, indicating that the numerous small puncta observed after 1 hour of induction coalesced into fewer, larger puncta as the cytoplasmic concentration increased over time. (Fig. 2C). However, cells overexpressing Nst1ΔPA and Nst1ΔPE failed to undergo coalescence, and showed an increased number of puncta and decreased signal intensities compared to cells overexpressing wildtype Nst1 and Nst1ΔPAM (Fig. 2C). Even after 24 hours of induction, wild-type Nst1 and Nst1ΔPAM formed mature condensates with intense fluorescence signals (Fig. 2C). However, cells expressing Nst1ΔPA and Nst1ΔPE did not show a reduction in puncta number, although some puncta increased in size and intensity compared to those observed after 4 hours of induction (Fig. 2C). This result indicates that PA domain is essential for coalescence of Nst1 condensates. Since several mutations in a protein can alter the physical property of condensates (Banani et al., 2022; Sanfeliu-Cerdan et al., 2023), we also determined whether these condensates exhibit liquid-like or amyloid-like characteristics. Cells overexpressing each domain deletion mutant for 4 hours were treated with 1,6-hexanediol for 1 hour, then observed. Most puncta generated by each mutant dispersed upon treatment, indicating that none of the domains significantly alters the liquid-like properties of these condensates (Fig. 2D).

To evaluate the condensation potential of each domain deletion mutant, we measured the protein concentrations associated with each intracellular puncta formation pattern for each Nst1 variant after 4 hours of induction by FCS (Fig. 2E). Cells expressing GFP or Nst1ΔAPD remained diffuse, whereas cells expressing Nst1ΔPA or Nst1ΔPE exhibited microphase separation without forming mature condensates, despite having cytoplasmic protein concentrations comparable to those of wild-type Nst1 and Nst1ΔPAM (Fig. 2E).

When we initially constructed the Nst1ΔPA mutant, positively charged amino acids near the C-terminal portion of the defined PA region were included in the deletion. To better distinguish the function of negatively charged clusters within this region, we generated an additional mutant called Nst1ΔPA-1 that excluded positively charged amino acids from deletion while maintaining the integrity of the negatively-charged domain within PA. Consistently, cells overexpressing Nst1ΔPA-1 displayed numerous micro-condensates from 1hour to 24 hours induction, similar to cells overexpressing Nst1ΔPA (Fig. S4). These findings indicate that the overexpression phenotype observed in Nst1ΔPA is due to the polyanion located within the PA region.

Next, we investigated whether the results from the Nst1 domain deletion mutants could be applied in condensate formation of endogenous Nst1. Since endogenously expressed Nst1 forms condensates from diauxic shift to stationary phase (Fig. 2F), we examined the puncta generation patterns by condensation of endogenously expressed full-length Nst1, the Nst1ΔPA, the Nst1ΔPAM, and the Nst1ΔAPD from early log phase to stationary phase (Fig. 2F).

We cultured cells expressing the chromosome-integrated GFP-tagged versions of each domain deletion mutant under the control of the endogenous Nst1 promoter in a *Δnst1* strain and observed them at 16, 30, 48, and 80 hours after seeding at an initial OD_600_ = 0.1 (Fig. 2G). These mutants exhibited similar puncta generation patterns to those observed in cells overexpressing these deletion mutants. Endogenously expressed Nst1ΔAPD formed few condensates, confirming that the aggregation-prone region, which is critical for hydrophobic core assembly, is essential for condensate formation (Fig. 2H-I). Among the tested mutants, Nst1ΔPA was the first to initiate condensation, starting at 10 hours during the early log phase (Fig. 2H-I), and notably during the mid-log phase (14–18 hours). While the wild-type Nst1 and other mutants had not yet begun puncta formation, Nst1ΔPA condensates increased significantly (Fig. 2H-I). After 18 hours, condensation began for the full-length Nst1 and Nst1ΔPAM. Interestingly, starting at 22 hours, as cells entered the stationary phase, the intensity of Nst1ΔPAM condensates were markedly higher than those of the wild-type Nst1 and Nst1ΔPA condensates (Fig. 2I). At 46 hours, during the stationary phase, we observed that the intensities of Nst1ΔPA condensates were lower than those of wild-type Nst1 and Nst1ΔPAM condensates (Fig. 2J). Together, these findings demonstrate that the differential propensities for condensate formation observed among mutants in overexpression experiments are also recapitulated under physiological conditions.

### Polyanion (PA) domain exclusively functions in coalescence of Nst1 condensates

Next, we investigated whether the contribution of the PA domain to Nst1 condensation is unique to the PA region, or whether other sequence regions of similar size can also play a similar role, thereby assessing potential non-specific effects arising from deletions of comparable regions. To clarify this, we generated five additional domain deletion mutants of similar size as the PA domain in Nst1: Nst1ΔD1^2-122^, Nst1ΔD2^123-293^, Nst1ΔD3^294-459^, Nst1ΔD4^753-844^, Nst1ΔD5^845-1015^ (Fig. 3A). The *Δnst1* w303a cells were transformed with plasmids that can overexpress each GFP-tagged Nst1 domain deletion mutant (D1∼D5). All strains were induced with galactose and observed over time using a fluorescence microscope at specified time points after the induction (Fig. 3B).

**Figure 3.**
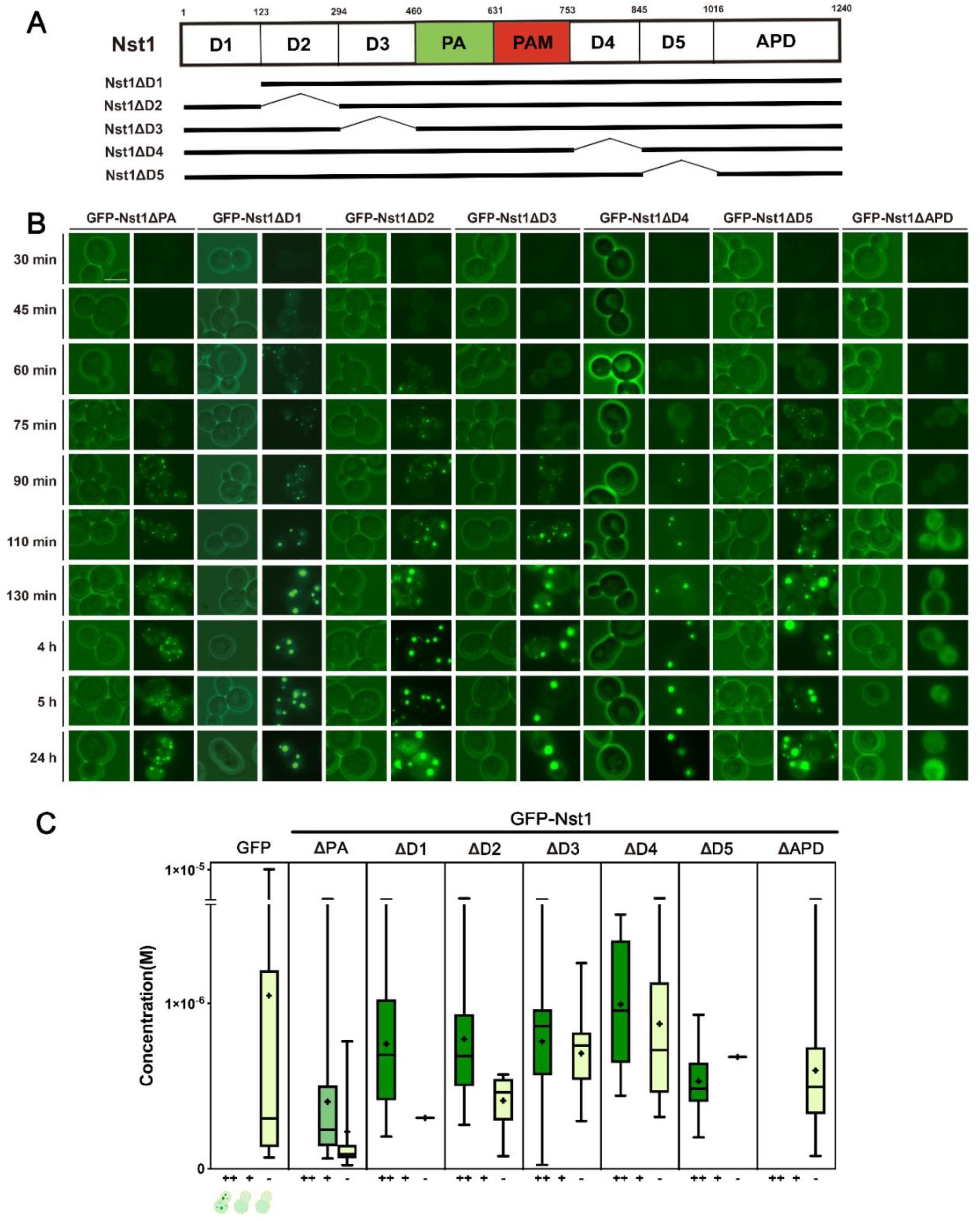
Polyanion (PA) domain exclusively functions in coalescence of Nst1 condensates. (A) A diagram of Nst1 with domain architectures. Residues 2-122 were designated as D1, 123-293 as D2, 294-459 as D3, 753-844 as D4, and 845-1015 as D5. The lengths of D1-D5 are similar to that of PA, and all of the domains are comparable in size. (B) Representative fluorescence microscopy images comparing time-dependent condensate formation in the PA deletion mutant and other domain deletion mutants (ΔD1-D5, ΔAPD). In *Δnst1* (YSK3490) cells, GFP-tagged Nst1ΔPA (pMW20-pGAL-GFP-NST1ΔPA), Nst1ΔD1 (pMW20-pGAL-GFP-NST1ΔD1), Nst1ΔD2 (pMW20-pGAL-GFP-NST1ΔD2), Nst1ΔD3 (pMW20-pGAL-GFP-NST1ΔD3), Nst1ΔD4 (pMW20-pGAL-GFP-NST1ΔD4), Nst1ΔD5 (pMW20-pGAL-GFP-NST1ΔD5), and Nst1ΔAPD (pMW20-pGAL-GFP-NST1ΔAPD) were overexpressed. Overexpression of each Nst1 derivative was induced by galactose, and images were obtained at the designated time point after the induction. Representative images were selected among more than 100 cells. Scale bar: 5 µm. (C) Box plot analysis showing the correlation between puncta formation patterns and cytoplasmic concentrations in the cells overexpressing GFP-tagged Nst1, Nst1ΔPA, NstΔD1-ΔD5, and Nst1ΔAPD after 4 hours of induction. Cytoplasmic protein concentrations were measured via fluorescence correlation spectroscopy (FCS). The puncta formation patterns in cells were classified into three categories: diffuse phase, microphase separation, and mature condensates.

Except the Nst1ΔAPD deletion mutant, the overexpression of D1∼D5 deletion mutants began to induce condensates approximately 1 hour after galactose induction (Fig. 3B). Similar to the Nst1ΔPAM (Fig. 2C, 3B), the condensates of the deletion mutants from D1 to D5, increased in size as the number of condensates per cell decreased, except for the Nst1ΔPA (Fig. 3B). When cytoplasmic protein concentrations were measured by FCS after 4 hours of induction, Nst1ΔPA exhibited microphase separation at all measured concentrations, whereas all other mutants formed mature condensates at similar cytoplasmic concentrations (Fig. 3C). Even after 24 hours of induction, Nst1ΔPA failed to generate mature condensates, while all the other mutants, ΔD1 to ΔD5, successfully formed mature condensates (Fig. 3B). These results further support that the PA region of Nst1 plays a unique and critical role in driving condensate coalescence during biomolecular condensation.

### The polyanion (PA) domain of Nst1 enhances intermolecular interaction within condensates, whereas the polyampholyte (PAM) domain of Nst1 increases the rigidity of Nst1 condensates

Next, we examined whether the PA and PAM domains influence the dynamics of Nst1 condensates by employing fluorescence recovery after photobleaching (FRAP). Since the condensates of the overexpressed Nst1ΔPA did not coalescence and retained minimal puncta size, the abnormal function in the coalescence process is likely related to the molecular dynamics affected by the PA domain deletion.

We examined GFP-tagged the full-length Nst1, as well as the Nst1ΔPA, Nst1ΔPAM, and Nst1ΔD1 mutants, with GFP control after 4 hours of induction (Fig. 4A). Condensates from each mutant were imaged, photobleached, and fluorescence recovery was monitored over time (Fig. 4B). Prior to be analyzed, recovery curves from each mutant FRAP analysis were fitted and normalized (Fig. 4C-D, S6 A-E). Because molecular dynamics can be affected by molecular size (Pack, 2021), the Nst1ΔD1 mutant was also used as a size control for the analyses of Nst1ΔPA and Nst1ΔPAM.

**Figure 4.**
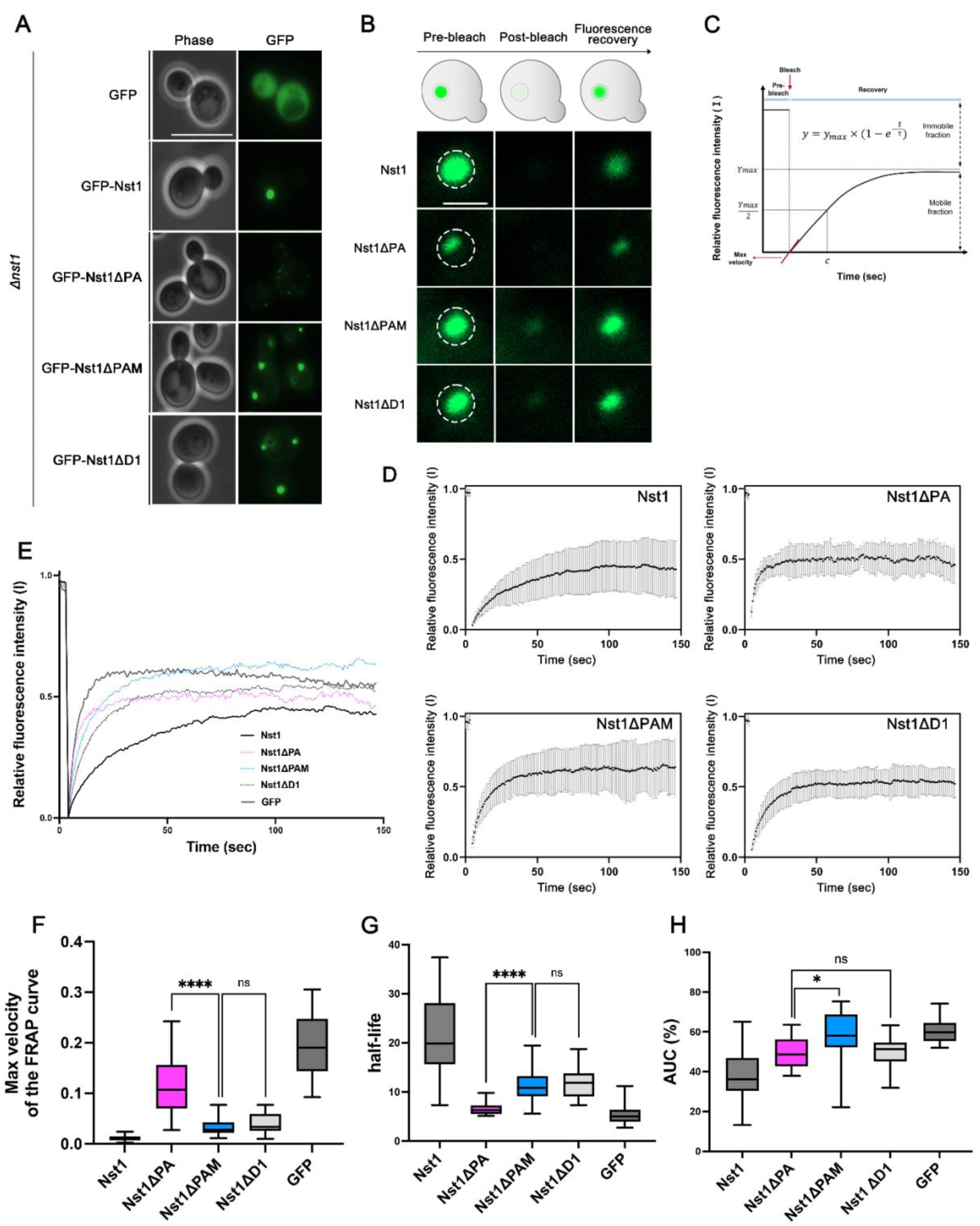
The polyanion (PA) domain of Nst1 enhances intermolecular interaction within condensates, whereas the polyampholyte (PAM) domain increases the rigidity of Nst1 condensates. (A-H) GAL promoter induction was performed for *Δnst1* (YSK3490) cells transformed with pMW20-pGAL - GFP-NST1, pMW20-pGAL-GFP-NST1ΔPA, pMW20-pGAL-GFP-NST1ΔPAM, pMW20-pGAL-GFP-NST1ΔD1. Fluorescence Recovery After Photobleaching (FRAP) experiments were performed on induced condensates of these cells as described in Materials and Methods. (A) Representative fluorescence microscopy images of cells overexpressing GFP-tagged full-length Nst1, Nst1ΔPA, Nst1ΔPAM, Nst1ΔD1, and GFP alone (control) in the *Δnst1* strain. Scale bar: 10 μm. (B) Representative fluorescence microscopy images at pre-beach point (cycle 1 of 200), post-bleach (cycle 6 of 200), and fluorescence recovery (cycle 200 of 200) during FRAP analysis. The stitched lines indicate the region of interest (ROI) selected for bleaching the puncta in cells. All experiments were performed with more than 20 puncta each. Scale bar: 1 μm. (C) Predicted FRAP recovery curve modeled as a single exponential function. 𝑡 is time, 𝑦_𝑚𝑎𝑥_ is the 𝑦 value at infinite time (𝑡 ≈ ∞), 𝜏 (tau) is the time point when the 𝑦 value is equal to 63.2% of 𝑦_𝑚𝑎𝑥_ value, and 𝑐 is half-life. (D) The average FRAP recovery curves of Nst1 and its mutants shown in (B). Each graph for full-length Nst1, Nst1ΔPA, Nst1ΔPAM, and Nst1ΔD1. The x-axis of these average graphs represents the time (sec) taken in 200 cycles, and the y-axis indicates the relative fluorescence intensity. (E) The average FRAP recovery curves for Nst1 and all Nst1 mutants in (D) were plotted together for direct comparison. Full-length Nst1 (black line), Nst1ΔPA (pink dotted line), Nst1ΔPAM (blue dotted line), Nst1ΔD1 (black dotted line), and GFP (grey line) as a control. The x-axis and y-axis represent the same values as in (D). (F) The maximum recovery rate of the FRAP curves for condensates of full-length Nst1, Nst1ΔPA, Nst1ΔPAM, and Nst1ΔD1. Akima spline interpolation was applied to all FRAP curves. For each Nst1 mutant, the first derivative (slope) at the point of maximal recovery rate was calculated using GraphPad Prism 10. The y-axis represents the maximum recovery rate (maximum slope) of the fitted FRAP curve. Statistical significance was determined using unpaired two-tailed Student’s t-test (**** p<0.0001) with Welch’s correction. P < 0.05 was considered statistically significant. (G) The half-lives of FRAP curves for condensates of full-length Nst1, Nst1ΔPA, Nst1ΔPAM, and Nst1ΔD1. The x-values corresponding to half-maximal recovery (y = 0.5) were calculated and averaged using GraphPad Prism 10. Statistical significance was determined using unpaired two-tailed Student’s t-test (**** p<0.0001) with Welch’s correction. P < 0.05 was considered statistically significant. (H) The area under the curve (AUC) of the FRAP recovery curves for condensates of full-length Nst1, Nst1ΔPA, Nst1ΔPAM, and Nst1ΔD1. The AUC was calculated for each normalized FRAP recovery curve using GraphPad Prism 10. Statistical significance was determined using unpaired two-tailed Student’s t-test (* p<0.01) with Welch’s correction. P < 0.05 was considered statistically significant.

We measured the maximal recovery velocity and recovery half-life for each Nst1 domain deletion mutant (Fig. 4C, F-G). The maximal recovery velocity represents the rate at which unbleached fluorescent molecules exchange with bleached molecules within puncta; a lower maximal recovery velocity indicates stronger intermolecular interactions within condensates, while a higher velocity suggests weaker interactions and increased molecular dynamics. The recovery half-life reflects the time required for fluorescence in the bleached region to reach half of its maximum equilibrium intensity, serving as an additional measure of molecular dynamics.

The full-length Nst1 exhibited the slowest and lowest maximal recovery velocity among all samples, whereas the PAM-, PA-, and D1-deleted mutants recovered more rapidly (Fig. 4D-F), indicating that molecules with smaller molecular weight are generally more dynamic in the condensates. Notably, Nst1ΔPA puncta showed the fastest maximal recovery velocity among all deletion mutants (Fig. 4F). In contrast, the maximal recovery velocity and half-life of Nst1ΔPAM were comparable to those of Nst1ΔD1 (Fig. 4F-G). These results suggest that Nst1ΔPA exhibits particularly increased molecular dynamics due to weaker and less stable intermolecular interactions compared to Nst1ΔPAM and Nst1ΔD1.

Regarding mobile fraction as area under curve (AUC, %), the full-length Nst1 had the lowest AUC compared with these Nst1 deletion mutant proteins (Fig. 4E). The AUC of the Nst1Δ PAM was significantly higher than those of the Nst1ΔPA and Nst1ΔD1 (Fig. 4H). This indicates that puncta formed by Nst1ΔPAM have a higher proportion of mobile fractions compared to those formed by Nst1ΔPA and Nst1ΔD1. This suggests that the PAM domain of Nst1 enhances the rigidity of Nst1 condensates.

Overall, our results strongly suggest that polyampholyte (PAM) deletion weakens cohesion of internal networks, increasing overall molecular mobility within the condensate, while the polyanion (PA) deletion specifically enhances single-molecule dynamics by reducing electrostatic molecular affinity. That is, polyanions might participate in intermolecular interactions, enforcing molecular affinity, while polyampholytes contribute to restricting the movement of molecules in the confined state within condensates. These results highlight how different charged domains contribute uniquely to condensate properties and molecular behavior.

### The PAM domain is an RNA-independent multivalent interaction domain for interaction with a P-body component, Dcp2

In our previous study, we found that overexpression of Nst1 induces the formation of Nst1 condensates, which simultaneously promote the condensation of other P-body components, including Dcp2 (Choi *et al*., 2022). We suggested that the PAM domain, previously described as PD, is a multivalent interaction region responsible for the condensation of P-body components such as Dcp2 (Choi *et al*., 2022).

Since the function of PAM domain was resolved by measuring endogenously expressing GFP-tagged Dcp2, Dhh1, and Xrn1 in cells overexpressing non-tagged Nst1 variants (Choi *et al*., 2022), we need to confirm the function of PAM domain using a direct approach that can elucidate its role in inducing Dcp2 condensation. We utilized the chromosomally mKate2-tagged Dcp2 strain overexpressing GFP-tagged Nst1 variants, including Nst1ΔPAM, Nst1ΔPA, and Nst1ΔD1. The cells were induced to generate condensates of each Nst1 variant by 4 hour-overexpression, then treated with cycloheximide which binds to the ribosomal A site and inhibits translation elongation, resulting in decrease of free RNA level (Schneider-Poetsch et al., 2010). The reduction of the cytoplasmic concentration of free RNA, thereby eliminates Dcp2 condensates that are formed by free RNA (Teixeira et al., 2005; Van Treeck and Parker, 2018). With this strategy, the majority of Dcp2 condensates formed by Nst1 variants could be observed under these conditions (Fig. 5A).

**Figure 5.**
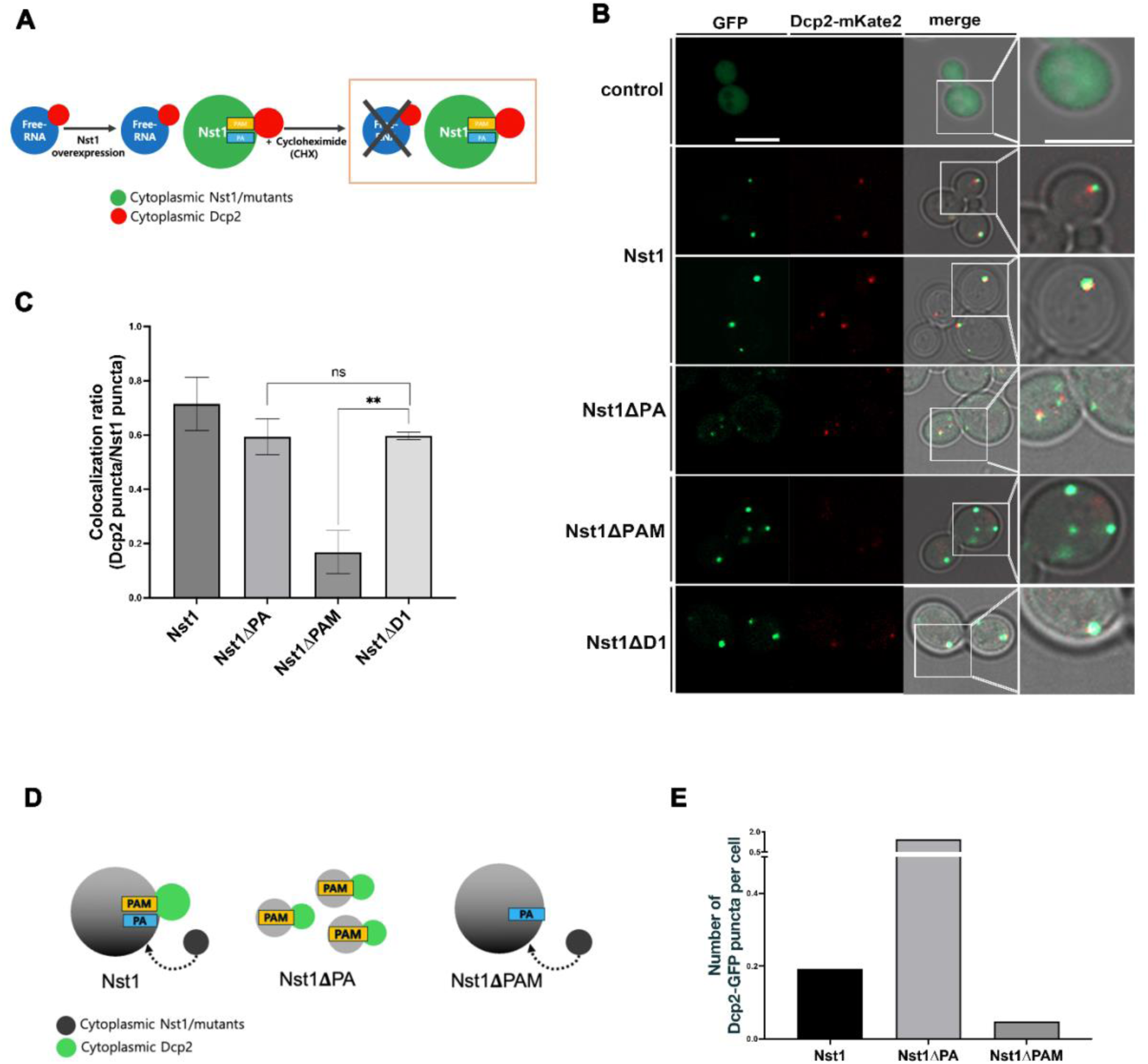
The PAM domain is an RNA-independent multivalent interaction domain for interaction with the P-body component Dcp2. (A) Schematic representation of the experiments (B-C) designed to examine the effect of RNA on P-body condensate formation of Nst1 and its PA and PAM deletion mutants. (B) High-resolution confocal microscopy images showing the formation of mKate2-tagged Dcp2 condensates induced by condensates of GFP-tagged Nst1, Nst1ΔPA, Nst1ΔPAM, and Nst1ΔD1. W303a *Δnst1* cells endogenously expressing Dcp2-mKate2 (YSK3663) were transformed with pMW20-pGAL-GFP-NST1, pMW20-pGAL-GFP-NST1ΔPA, pMW20-pGAL-GFP-NST1ΔPAM, or vector only (control). Following galactose induction for 4 hours, cells were treated with cycloheximide (CHX, 100 μg/ml) for 10 minutes and imaged using a Zeiss LSM 980 confocal microscope equipped with an Airyscan 2 detector (Carl Zeiss). The right panel shows a 2× magnified view of selected puncta in the merged images. GFP and mKate2 channels are shown in green and red, respectively. Scale bar: 5 µm. Representative images are shown. (C) Quantitative analysis of the spatial co-localization between Dcp2 and Nst1 condensates shown in (B) after CHX-treatment. The co-localization ratio was calculated as the proportion of Dcp2 condensates overlapping with Nst1 and its mutants-derived puncta. Puncta were segmented and quantified using Fiji (ImageJ) software. For each Nst1 and its mutants, at least 100 condensates were analyzed per experiment across three independent replicates. Statistical significance was determined by unpaired t-test (***p < 0.001) using GraphPad Prism 10. (D) Schematic representation of the proposed Dcp2 condensation model based on the functional roles of the PA and PAM domains of Nst1. (E) Quantification of Dcp2-EGFP puncta formation in yeast cells (YSK3709) imaged by widefield epifluorescence microscopy. Cells endogenously expressing Dcp2-EGFP were transformed to overexpress Nst1, Nst1ΔPA, or Nst1ΔPAM for 4 hours. Images were acquired using identical microscope settings for all conditions. Dcp2-EGFP puncta were segmented in FIJI (ImageJ) by intensity thresholding. The number of puncta per cell was quantified and normalized by cell number, with more than 300 cells analyzed for each condition. Data are presented as bar graphs generated using GraphPad Prism 10.

Using high-resolution confocal microscopy, we clearly observed the formation of multiphasic condensates composed of Nst1 and Dcp2 (Fig. 5B). Most wild-type Nst1 condensates were co-localized with mKate2-tagged Dcp2, suggesting that Nst1 condensates promote the formation of Dcp2 condensates (Fig. 5C). Interestingly, the fluorescence signals from Nst1 and Dcp2 condensates did not completely overlap; instead, the two signals appeared as if distinct condensates were docked together (Fig. 5B).

Both Nst1ΔPA and Nst1ΔD1 condensates also exhibited co-localization with Dcp2-mKate2 condensates (Fig. 5B-C). Although Nst1ΔPA condensates were able to induce Dcp2 condensation, their size was relatively smaller compared to other Nst1 variant condensates, consistent with the unique role of PA in coalescence of Nst1’s microphase condensates (Fig. 5B). In contrast, Nst1ΔPAM condensates rarely promoted the formation of Dcp2 condensates, highlighting the critical role of the PAM domain in facilitating Dcp2 condensation upon Nst1 overexpression (Fig. 5B). Condensates formed by overexpression of GFP-tagged Nst1 and its mutants demonstrated no change in the integrated density of puncta before and after CHX treatment (S7. A). This indicates that a decrease in cytoplasmic concentration of ribosomal free RNA induced by CHX treatment does not affect Nst1 condensation.

To confirm the role of PAM on Dcp2 condensation, we employed an additional experimental approach to quantitatively measure Dcp2 condensation induced by overexpression of Nst1ΔPA and Nst1ΔPAM in each cell. If the PAM domain induces Dcp2 condensation directly, rather than via RNA, then the number of Dcp2 condensates should not be reduced in Nst1ΔPA cells even after cycloheximide treatment, because Nst1ΔPA condensates, which contain the PAM domain, fail to coalesce and instead increase in number. In contrast, Nst1ΔPAM should lead to a significant reduction in the number of Dcp2 condensates (Fig. 5D). For the quantitative analysis of Dcp2 condensates induced by condensation of wild-type Nst1 and its domain deletion mutants, instead of relying on the weak signals from mKate2-tagged Dcp2, we employed a GFP-tagged Dcp2 strain that allowed more robust quantification to further validate the function of the PAM domain (Fig. 5D). Cells overexpressing either non-tagged full-length Nst1 or the deletion mutants (Nst1ΔPA, Nst1ΔPAM) for 4 hours were treated with cycloheximide for 10 minutes prior to imaging. Subsequently, the puncta formed by chromosomally EGFP-tagged Dcp2 in these overexpression conditions were segmented and quantified (Fig. 5D-E). Notably, cells overexpressing Nst1ΔPA exhibited a substantially higher number of Dcp2 puncta per cell compared to cells overexpressing wild-type Nst1 (Fig. 5E), suggesting that the increased number of Nst1ΔPA condensates containing the PAM domain may promote the formation of a greater number of Dcp2 condensates than wild-type Nst1. In contrast, the number of Dcp2 puncta per cell was reduced in Nst1ΔPAM-overexpressing cells relative to the wild-type (Fig. 5E), indicating that the PAM domain provides a physical platform necessary for efficient Dcp2 condensation.

### The Polyanion (PA) domain of human TDP-43 promotes coalescence of condensates, similar to Nst1

We expanded the functional analysis of PA and PAM subdomains to the scaffold protein Edc3 focusing on their roles in biomolecular condensation. Since Edc3 also has a distinctive negative charged amino acid cluster in PE domain, we divided the PE domain of Edc3 into two parts: the PA region, characterized by a cluster of negatively charged amino acids, and the PAM region, where positively and negatively charged amino acids are more evenly distributed (Fig. S2, S8). We constructed plasmids for the overexpression of GFP-tagged Edc3ΔPAM and Edc3ΔPA mutants, and overexpressed them together with GFP-tagged Edc3, Edc3ΔPE, and Edc3ΔAPD in the *Δedc3* strain to examine their condensate phenotypes. Remarkably, as observed with Nst1, after 4 hours of overexpression, Edc3 formed mature condensates, whereas Edc3ΔPA, similar to Edc3ΔPE, exhibited microphase separation in all cells (Fig. S8). Even after 24 hours of induction, Edc3ΔPA failed to form mature condensates in almost all cells, and only microphase-separated condensates were observed, demonstrating the role of PA in coalescence of Edc3 condensates. Edc3ΔAPD, consistent with the result of Nst1, did not form condensates (Fig. S8).

We then investigated whether the phenotype of PA deletion mutants observed in the budding yeast proteins Nst1 and Edc3 is also present in proteins with self-condensation potential that are implicated in neurodegenerative diseases. Since TDP-43 are known to undergo self-condensation upon overexpression in budding yeast (Fushimi et al., 2011; Johnson et al., 2008; Outeiro and Lindquist, 2003; Sun et al., 2011), we first confirmed TDP-43 aggregation upon overexpression in budding yeast as been previously reported. Under the same overexpression conditions used for Nst1 and Edc3, TDP-43 also underwent self-condensation in budding yeast (Fig. 6B). Two PA domains (PA1, PA2) in the TDP-43 sequence were designated with using the sliding window approach (Fig. 6A, S3). The low complexity region in the C-terminal domain (CTD) has been known to be critical for TDP-43 amyloid aggregation (Chien et al., 2021; Li et al., 2021; Shenoy et al., 2023). The CTD has a relatively low charged amino acid cluster score, but within the CTD, residues 255-276, which have a relatively high density of charged amino acid, were designated as the PAM domain. To identify intrinsic sequence determinants that consists of charged amino acid cluster of TDP-43 aggregation, we generated and overexpressed those deletion mutants of TDP-43 in the budding yeast system and monitored them over time. Strikingly, the PA1 and PA2 deletion mutants failed to form mature condensates at 4 hours of overexpression compared to the full-length TDP-43, suggesting their roles in coalescence of condensates as observed role of PA in Nst1 and Edc3 (Fig. 6B).

**Figure 6.**
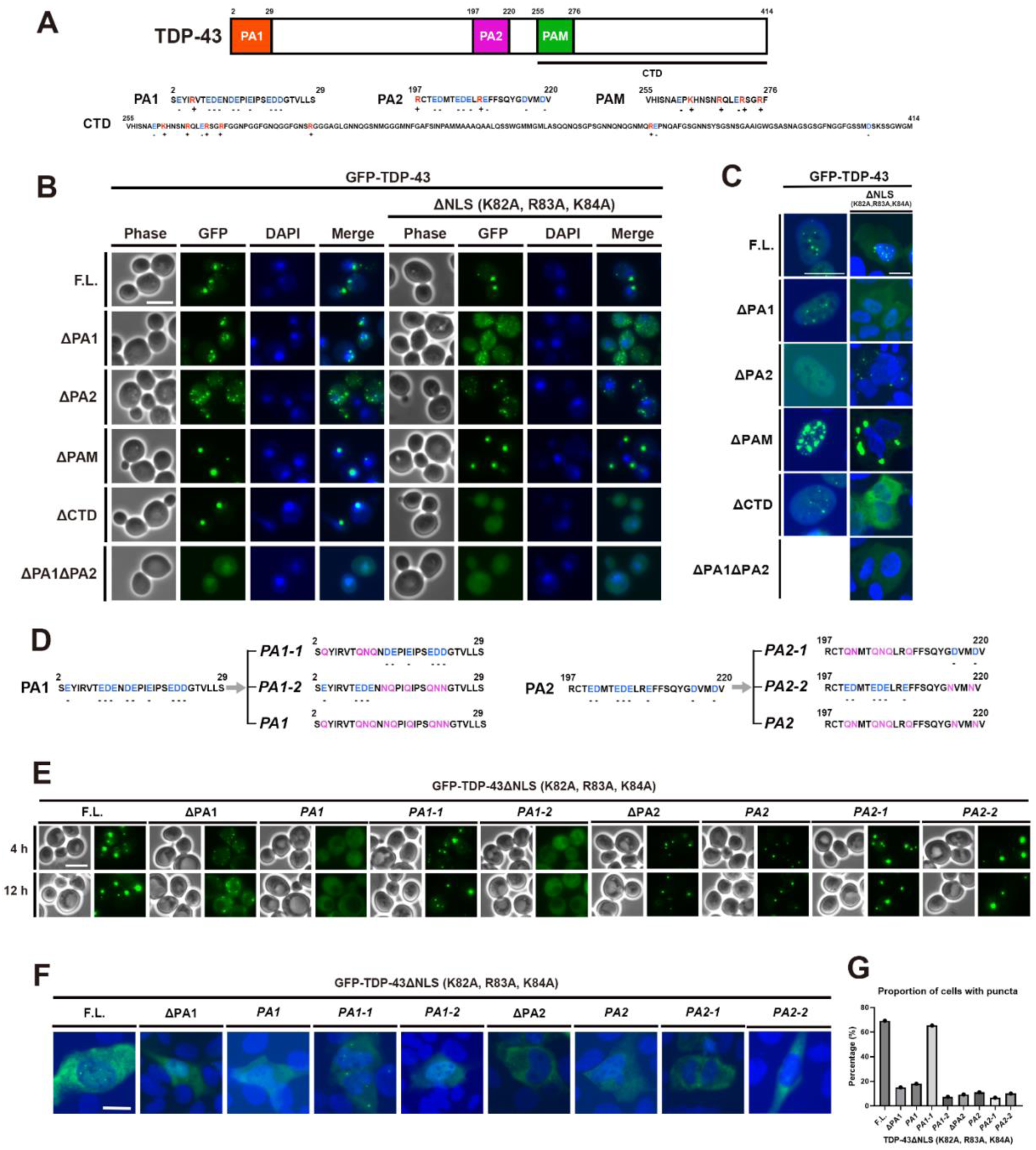
The Polyanion (PA) domain of Edc3 and human TDP-43 promote coalescence of condensates, similar to Nst1. (A) Schematic representation of TDP-43 domain architecture. Residues 2–29 and 197–220 were identified as clusters of negatively charged amino acids, designated PA1 and PA2, based on a sliding window approach and prediction tools, as described in Supplementary Figure 3. The region spanning residue 255-276 was denoted as PAM, while residues 255-414 were characterized as the C-terminal domain. See also Supplementary figure 3. (B) Representative widefield fluorescence microscopy images of budding yeast (W303a) cells after 4 hours of overexpression of TDP-43 and its domain deletion mutants (ΔPA1, ΔPA2, ΔPAM, ΔCTD and ΔPA1ΔPA2) with or without NLS (Nuclear localization signal) sequence under a galactose-inducible promoter. Cells were fixed and stained with DAPI to visualize nuclei. Scale bar: 5 µm. (C) Representative wildfield fluorescence microscopy images of cells expressing GFP-tagged TDP-43 and its variants (ΔPA1, ΔPA2, ΔPAM, ΔCTD and ΔPA1ΔPA2) in U2OS cells at 48 hours after transfection. U2OS cells were transfected with plasmids encoding GFP-tagged TDP-43 variants under CMV promoter as described in Materials and Methods. After 48 hours, cells were observed using fluorescence microscopy to assess condensation pattern of each TDP-43 variant. Scale bar: 20 μm. (D) The amino acid sequence of the PA1 and PA2 domains of TDP-43, along with its *PA1-1, PA1-2, PA1, PA2-1, PA2-2,* and *PA2* mutant variants, is depicted at the top. In the *PA1-1, PA1-2, PA1, PA2-1, PA2-2,* and *PA2* mutants, the negatively charged glutamic acid (E) and aspartic acid (D) residues were individually substituted with glutamine (Q) and asparagine (N), respectively, as indicated. (E) Representative widefield fluorescence microscopy images are shown for budding yeast (W303a) cells overexpressing ΔNLS TDP-43 full-length (F.L.), each PA domain deletion mutants (ΔPA1 and ΔPA2), as well as the *PA1, PA1-1, PA1-2, PA2, PA2-1*, and *PA2-2* variants, all of which lack the NLS. Images were taken following 4 and 12 hours of galactose induction. For each strain and time point, more than 100 cells were examined, and representative images are presented. Scale bar: 5 μm. (F) Representative fluorescence microscopy images of U2OS cells expressing GFP-tagged ΔNLS TDP-43 and its variants (ΔPA1, *PA1*, *PA1-1*, *PA1-2*, ΔPA2, *PA2*, *PA2-1* and *PA2-2*) 48 hours post-transfection. U2OS cells were transfected with plasmids encoding each GFP-tagged ΔNLS TDP-43 construct under the control of the CMV promoter, as detailed in the Materials and Methods. After 48 hours, cells were imaged by fluorescence microscopy to evaluate the condensation behavior of each variant. (G) Bar plot showing the percentage of cells containing TDP-43 puncta in U2OS cells expressing TDP-43 lacking the NLS and its charge-removal mutants. Scale bar, 20 μm.

The previously characterized CTD deletion mutant having NLS sequence also formed condensates when overexpressed, similar to the reported study (Johnson *et al*., 2008; Wang et al., 2018a). The morphology of these condensates differed from that of the full-length TDP-43: while the full-length TDP-43 formed both large and small, irregularly shaped condensates, the CTD deletion mutant produced very large, round condensates in much lower numbers per cell (Fig. 6B). Interestingly, the condensate morphology of the TDP-43ΔCTD mutant was recapitulated in the TDP-43ΔPAM mutant in the nucleus, suggesting that the phenotype in the nucleus might be attributable to the loss of the PAM domain within the CTD rather than the entire NTD existence or CTD deletion. This implies that the charged amino acid cluster within the CTD may play a key role in determining the physical properties and dynamics of TDP-43 condensates in the nucleus. Since these TDP-43 mutants generated retain the nuclear localization signal (NLS), all overexpressed mutants predominantly localize to and form condensates in the nucleus. To observe the condensates of mutants in the cytoplasm, we additionally generated NLS-mutated TDP-43 variants. These variants exhibited phenotypes similar to those observed in the nucleus, except for the CTD-deleted TDP-43 variant: the PA1 and PA2 deletion mutants failed to form mature condensates (Fig 6B). Notably, while overexpressed TDP-43ΔCTD forms condensates in the nucleus, it remains diffusely distributed in the cytoplasm (Fig. 6B). The phenotypic pattern observed for each mutant after 4 hours of induction remained unchanged in cells induced for 8, 10, or 12 hours (S9).

To determine whether the condensate phenotypes of PA1, PA2, PAM, and CTD deletion mutants of TDP-43 observed in budding yeast are also present in mammalian cells, we overexpressed GFP-fused TDP-43 and TDP-43 mutants in U2OS cells. Consistent with our findings in budding yeast, the PA1 and PA2 deletion mutants formed less mature condensates in the nucleus, compared to the wildtype TDP-43 (Fig. 6C). In particular, the PA2 deletion mutant rarely formed condensates in the nucleus among all cells expressing the protein, as indicated by the presence of GFP signal (Fig. 6C).

To observe the condensation of TDP-43 derivatives in the cytoplasm, we introduced an additional NLS mutation (K82A, R83A, K84A) into each GFP-fused TDP-43 domain deletion mutant and examined their expression in U2OS cells (Lu et al., 2022). The cytoplasmic expression of these TDP-43 derivatives displayed a similar pattern of puncta formation to their nuclear counterparts. In the case of the ΔPA1ΔPA2 double deletion, condensate formation was further inhibited. These data confirm that PA domains are a key contributor to condensation of TDP-43. Notably, the PAM domain deletion mutant exhibited a markedly larger size and higher fluorescence intensity of puncta compared to other mutants both in nucleus and cytoplasm (Fig. 6C).

We next investigated whether the TDP-43 coalescence tendency observed for the PA1 and PA2 mutants was attributable to the negative charges of the clustered amino acids. To this end, we generated *PA1-1*, *PA1-2*, *PA2-1*, and *PA2-2* mutants in which the glutamate (E) and aspartate (D) residues in the partial N-and C-terminal regions of PA1 and PA2 were individually substituted with glutamine (Q) and asparagine (N), respectively, as well as *PA1* and *PA2* mutants in which all glutamate and aspartate residues were replaced with glutamine and asparagine (Fig. 6D).

Upon galactose induction in budding yeast cells, the PA1 and PA2 mutants with all negatively charged amino acids replaced by uncharged glutamine and asparagine exhibited a completely different profile in puncta formation compared to the wildtype TDP-43 protein (Fig. 6E). Unlike the ΔPA1 mutant, the *PA1* mutant exhibited a distinct phenotype: while the overexpressed ΔPA1 mutant displayed microphase separation, the overexpressed *PA1* showed a pronounced inhibition of condensate seeding (Fig. 6E). The *PA1-1* mutant, with partial N-terminal substitution, exhibited morphology identical to wildtype condensates, whereas *PA1-2* which includes E17Q and E21Q, resembled the *PA1* mutant, suggesting consistency with previous findings that E17 and E21 in the TDP-43 N-terminal domain are critical residues required for oligomerization (Afroz et al., 2017).

Notably, the results for the PA2 domain are remarkable. The *PA2* exhibited condensate morphology identical to that of ΔPA2 and displayed a pronounced tendency toward microphase separation, unlike the wildtype TDP-43 which forms mature condensates, (Fig. 6D). Unlike the *PA1-1* and *PA1-2* mutants, both the *PA2-1,* the *PA2-2* mutant which have partial substitutions of negatively charged amino acids in either the N-terminal or C-terminal regions failed to fully recapitulate the puncta generation patterns of ΔPA2 (Fig. 6D). These results indicate that it is not specific negatively charged residues within PA2 that are critical, but rather that the overall clusters of negatively charged amino acids in PA2 contribute to TDP-43 coalescence.

We introduced the same amino acid substitution mutants into U2OS cells. Microscopic images were captured for each mutant, and the portion of puncta-forming cells was quantified across the GFP-expressing cell population (Fig. 6F–G). While the TDP-43 mutant lacking the NLS but harboring no additional mutations formed small yet distinct cytoplasmic puncta, puncta formation was markedly inhibited in all substitution mutants with the exception of *PA1-1* (Fig. 6F–G). These results suggest the possibility that negative charges within the negatively charged amino acid clusters serve as critical regulators of TDP-43 phase behavior.

## DISCUSSION

Protein condensation in cell is not a binary switch between soluble and demixed states, but rather a multi-step process involving distinct kinetic phases: evolving from dynamic droplets into more stable, gel-like, or solid states within the cellular environment (Shen et al., 2023; Wang et al., 2021). The maturation of condensates is essentially driven by thermodynamic coarsening mechanisms—such as coalescence and Ostwald ripening—and is not merely a physical hardening but a biochemical cascade that facilitates pathological transitions(Banani *et al*., 2017). By concentrating condensation-prone proteins through these coarsening dynamics, condensates lower the energetic barrier for amyloidogenesis, creating a fertile environment where initial oligomeric species can form and act as seeds to template the conversion of native proteins into amyloid fibrils (Berry et al., 2015; Dickson and Golding, 2022; Sim and Creamer, 2002; Visser *et al*., 2024). LCRs are closely associated with phenotypic variation (Fondon and Garner, 2004) and function as multivalent interaction hubs, playing a critical role in protein condensation and amyloidogenesis (Banjade and Rosen, 2014; Linsenmeier *et al*., 2023). Collectively, these findings strongly suggest that LCRs are physiologically and pathologically important components in complex biological systems.

Charged amino acid clusters within LCRs represent one of the most frequently observed sequence motifs within proteins consisting of RNP granules (Niedner-Boblenz et al., 2024; Regy et al., 2020; Saitoh et al., 2004). Within intrinsically disordered regions (IDRs), there have been numerous clues that certain charged amino acids may contribute to the regulation of condensation, either by promoting or inhibiting condensation (Saar et al., 2021). Both negatively and positively charged amino acids are thought to modulate intermolecular affinity via electrostatic interactions, where the pattern of charged residues along the sequence plays a pivotal role. As mentioned previously, when charged residues are uniformly distributed, the driving force for phase separation is attenuated; in contrast, when charged residues are clustered or segregated from other residues, the propensity for phase separation is enhanced (Borcherds et al., 2021; Saitoh *et al*., 2004; Wang *et al*., 2018b). This study presents the first report on the differential roles of low-complexity regions (LCRs) composed of charged amino acids as sequence elements specifically regulating phase of the condensates.

With budding yeast proteins, Nst1 and Edc3, we demonstrate that the specific deletion and mutation of the negatively charged amino acid clusters termed polyanion (PA) domain in Nst1 and Edc3 lead to a distinct defect in coalescence, without abolishing the initial nucleation capability. This implies that the grammar governing assembly is distinct from the grammar governing fluidity and fusion. While multivalent interactions via PAM domains or RNA binding may drive nucleation (Van Treeck and Parker, 2018), the negative charged amino acid clusters in Nst1 and Edc3 appear to be a prerequisite for this maturation trajectory and act as ‘cohesion-tuners’ that enable the fusion between microphase condensates. FRAP analysis of Nst1 provides a mechanistic explanation for these phenotypes. The significantly rapid FRAP recovery of the Nst1ΔPA mutant suggests that the failure to coalesce is due to a loss of cohesive force (surface tension) rather than rigidification. Specifically, the observed “hyper-dynamic but non-coalescing” state suggests that this is not merely due to a loss of molecular stickiness, but rather because the interfacial tension between the droplets and the surrounding cytoplasm is reduced to a point where there is no energetic favorability to drive Ostwald ripening or coalescence (Brangwynne et al., 2009).

Another important aspect of this study is that this “coalescence grammar” is conserved in the human disease protein TDP-43. While both the ΔPA1 and ΔPA2 mutants exhibited coalescence-inhibited condensates, their corresponding negative charge substitution mutants diverged significantly. The PA1 led to a complete loss of condensates, a phenotype faithfully reproduced by the PA1-2 mutant, which completely suppressed condensate nucleation. This result corroborates previous findings that residues E17 and E21 are critical for TDP-43 oligomerization(Afroz *et al*., 2017; Wang *et al*., 2018a). In contrast, the *PA2* displayed a phenotype of impaired condensate coarsening, which was identical to that of the ΔPA2. Furthermore, the observation that neither the *PA2-1* nor the *PA2-2* partial substitution mutants could fully recapitulate the complete *PA2* phenotype suggests that the entire negative charge cluster within the PA2 region collectively functions in regulating condensate coarsening.

Taken together, these findings imply that the functional role of a negatively charged amino acid residue in the condensation process is highly context-dependent. This underscores the necessity for a more nuanced investigation into the diverse functions and positional effects of negative charge clusters in the regulation of protein condensation. Although we did not directly measure the biophysical properties of TDP-43 mutants, the phenotypic similarity strongly suggests a conserved mechanism: these negatively charged residues likely provide the necessary cohesive potential to form and mature TDP-43 condensates in cells (Afroz *et al*., 2017; Conicella et al., 2016). Thus, the negative charge clusters are essential for tuning the phase separation threshold and condensate properties, presumably acting as a molecular rheostat that controls whether TDP-43 achieves sufficient condensate maturation to propagate pathological states or remains in a more soluble, dispersed form.

A remarkable region within TDP-43 is the PAM domain, which appears to play a critical role in regulating condensate dynamics. In U2OS cells, we observed an unexpected phenotype: the deletion of the PAM domain—regardless of NLS mutation status—resulted in significantly enlarged condensates. This phenotype, which was not evident in budding yeast, implies that the PAM domain functions to inhibit condensate coarsening, standing in direct contrast to the PA domain.

Notably, the PAM domain encompasses Lysine 263 (K263), a residue critical for RNA interaction. The K263E mutation has been identified as a pathogenic variant that drastically reduces RNA binding affinity by disrupting electrostatic interactions with the RNA backbone, leading to severe neurotoxicity (Austin et al., 2014; Imaizumi et al., 2022). Paradoxically, this loss of binding capacity is accompanied by increased thermal stability, rendering the protein structurally rigid and prone to aggregation (Austin *et al*., 2014; Chen et al., 2019). These findings suggest that the enlarged condensates observed in PAM-deleted TDP-43 are presumably driven by a similar mechanism: a reduction in RNA binding affinity. Therefore, a detailed investigation into the role of charged amino acids within the PAM domain is essential to fully understand the molecular mechanisms underlying these observations. Given its pivotal role in preventing aberrant condensate growth, the PAM domain offers a critical foundation for elucidating TDP-43 phase separation dynamics and represents a promising target for future therapeutic investigations.

Finally, our study underscores the indispensability of investigating condensate physics directly within the cellular environment. The distinction between a “matured condensates” and a “hyper-dynamic, non-coalescing seed”, as observed in Nst1, is impossible to discern using static imaging or pure in vitro systems lacking cellular crowding and energy (Snead and Gladfelter, 2019). By employing live-cell biophysical tools, we established a more physiologically relevant framework for understanding condensate dynamics. This approach suggests that future therapeutic strategies should focus on modulating these specific sequence motifs to restore the material properties of condensates within the living cell.

## STAR★METHODS

### KEY RESOURCES TABLE

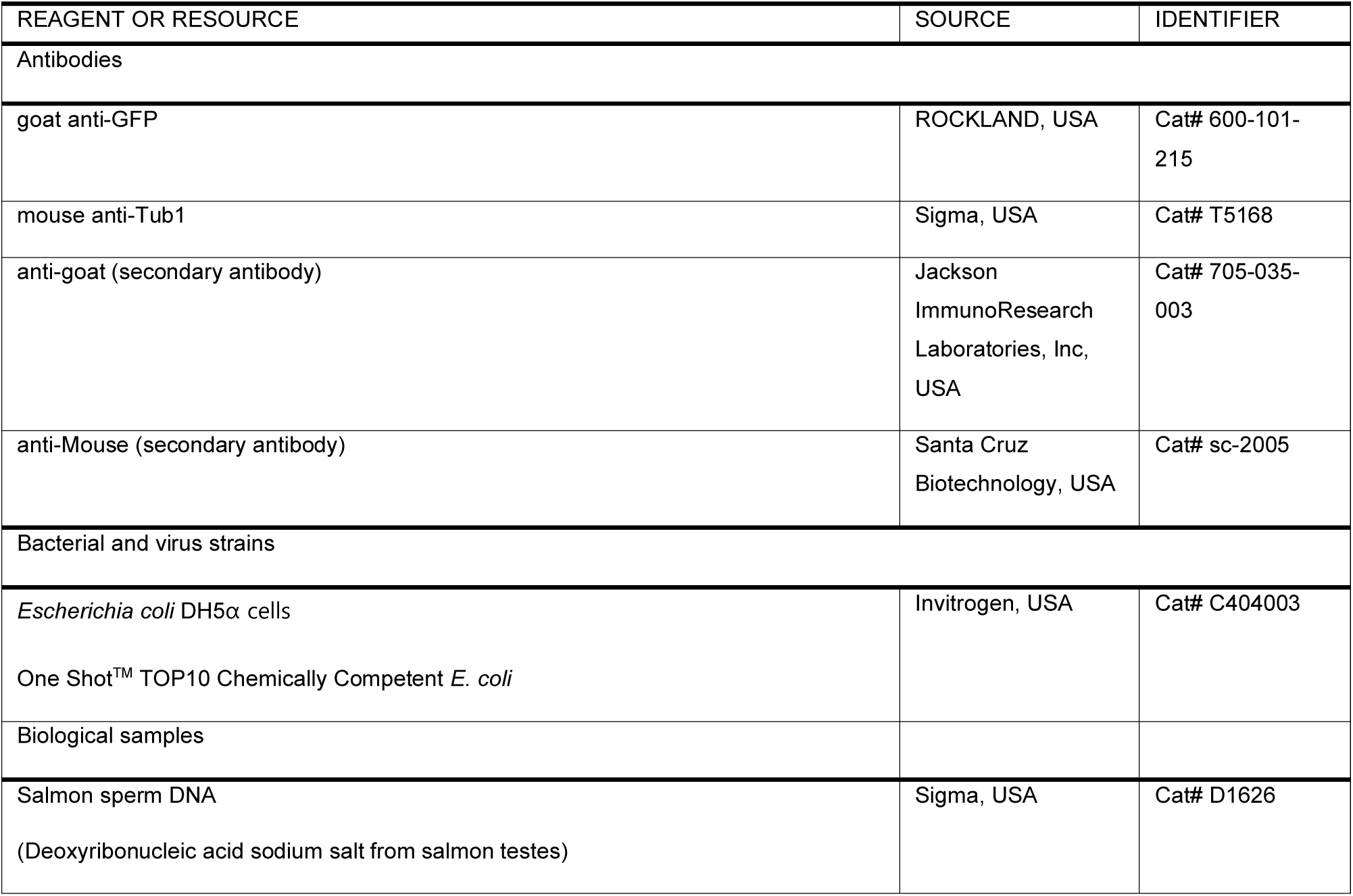
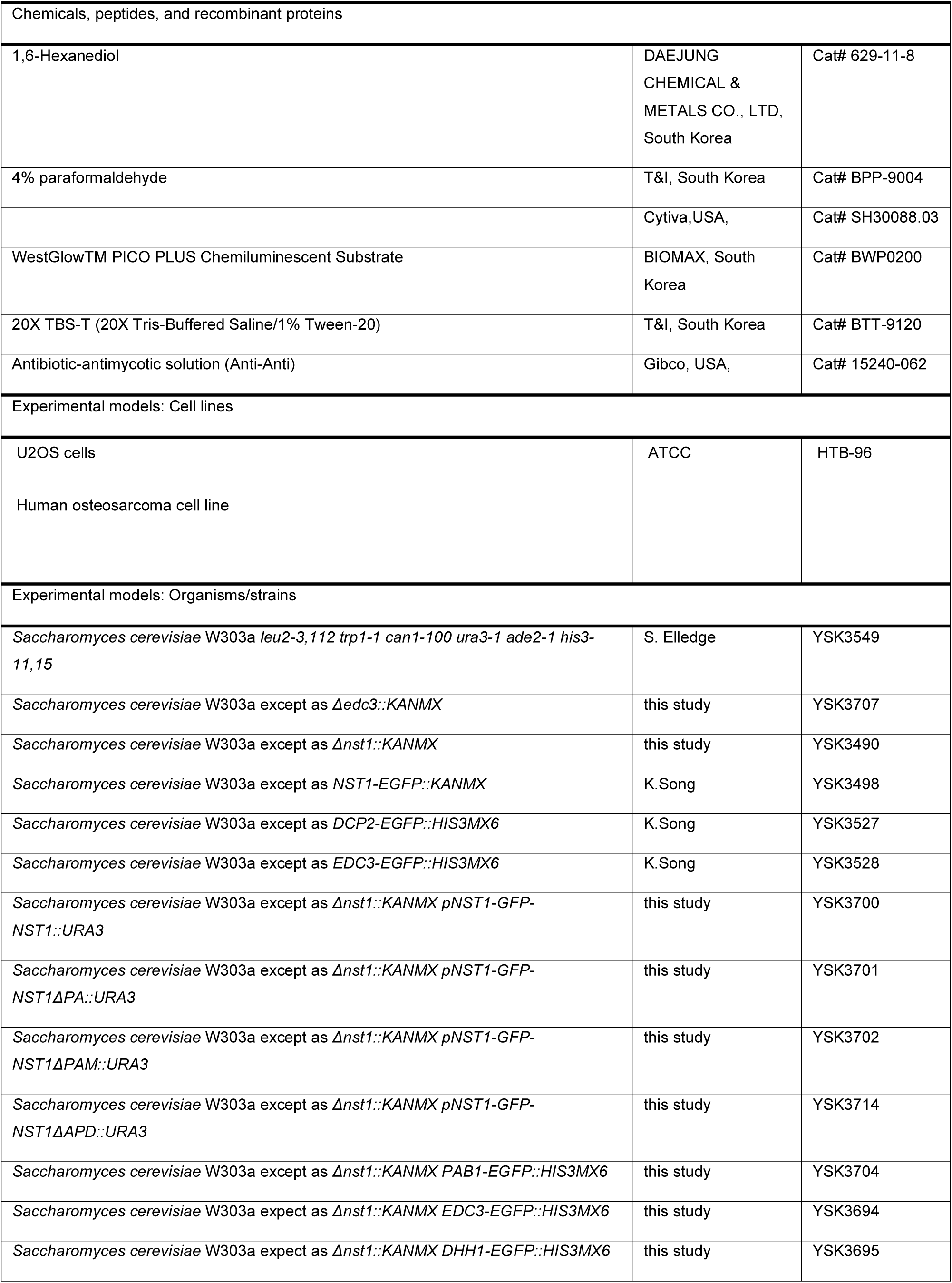
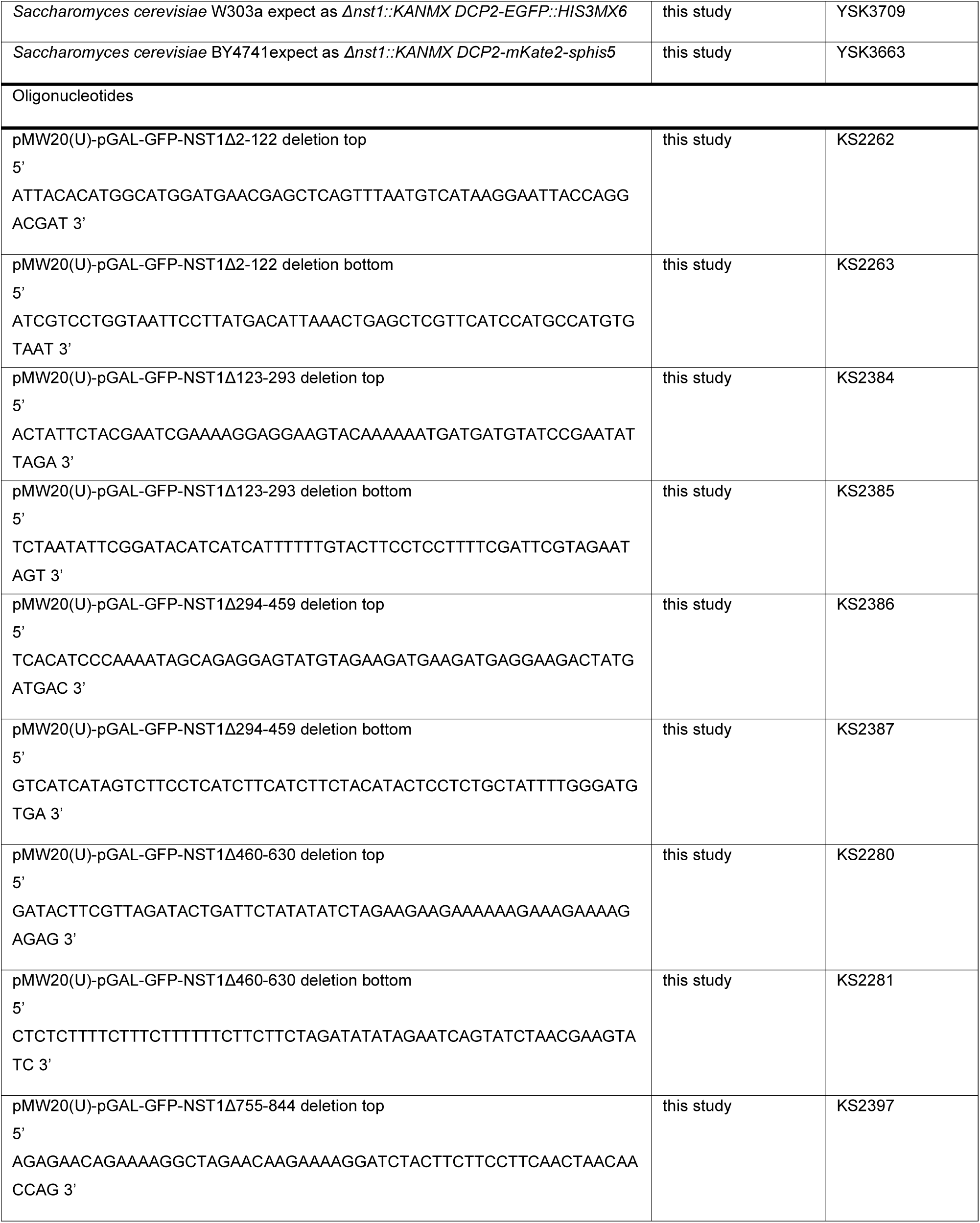
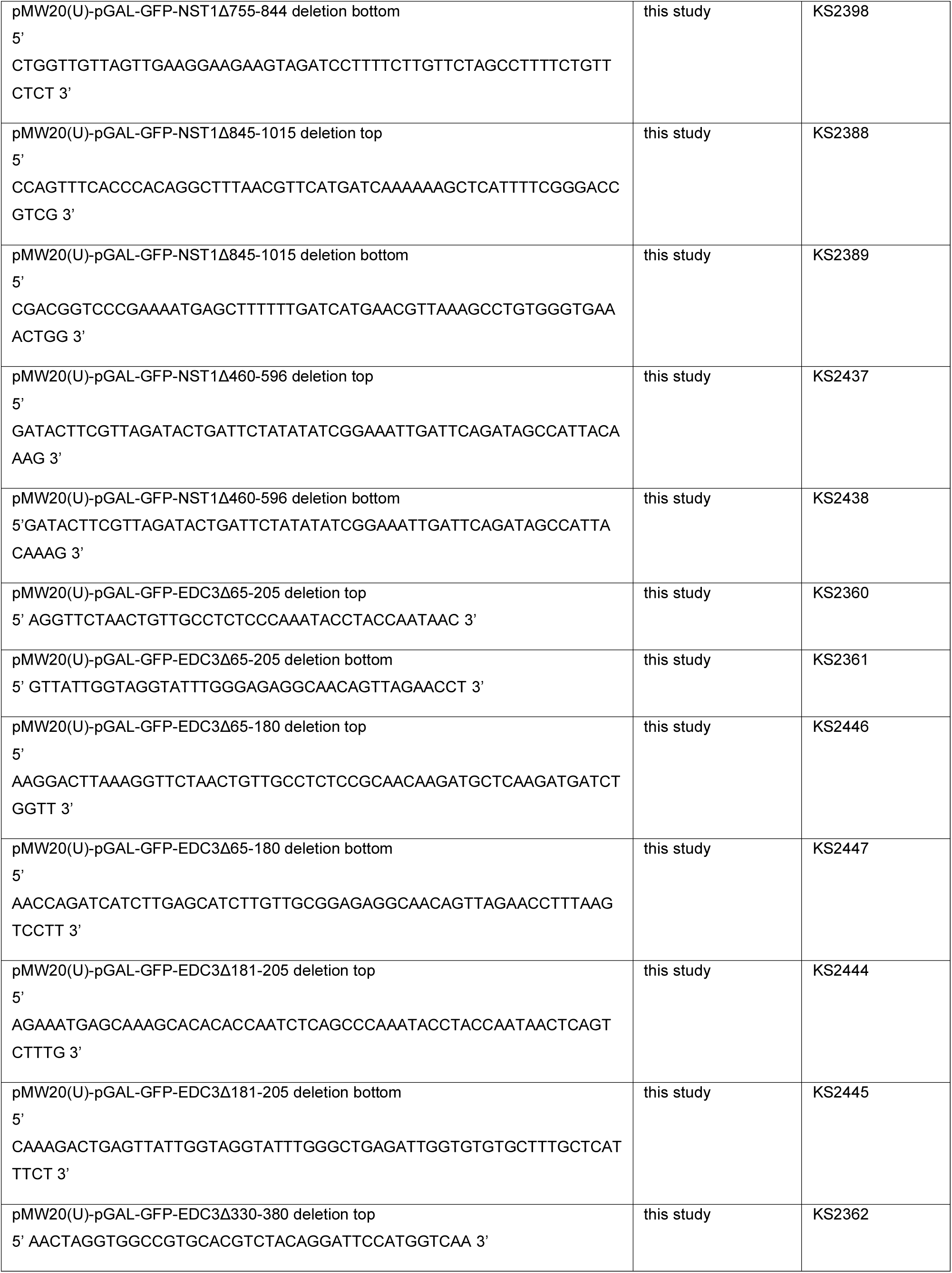
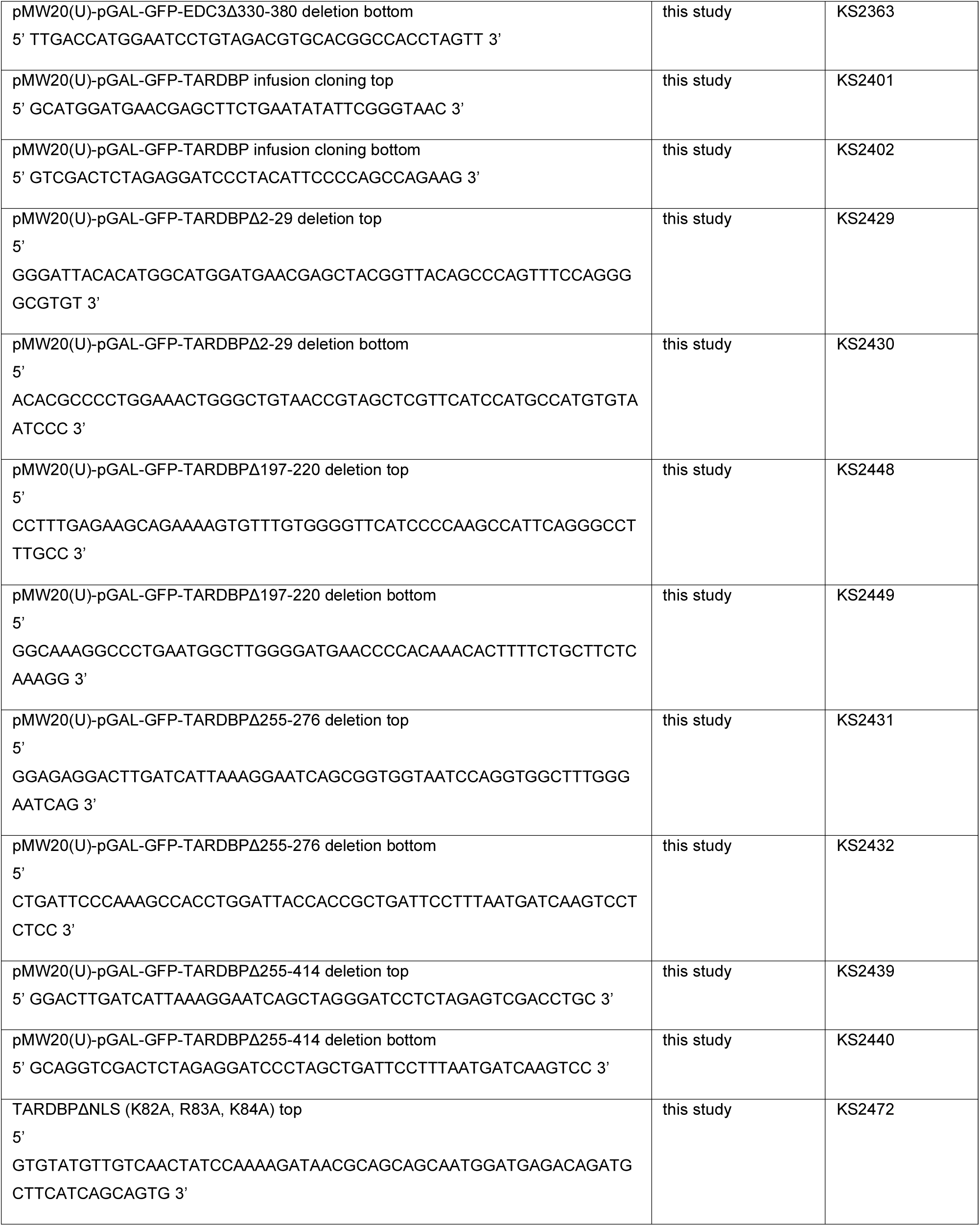
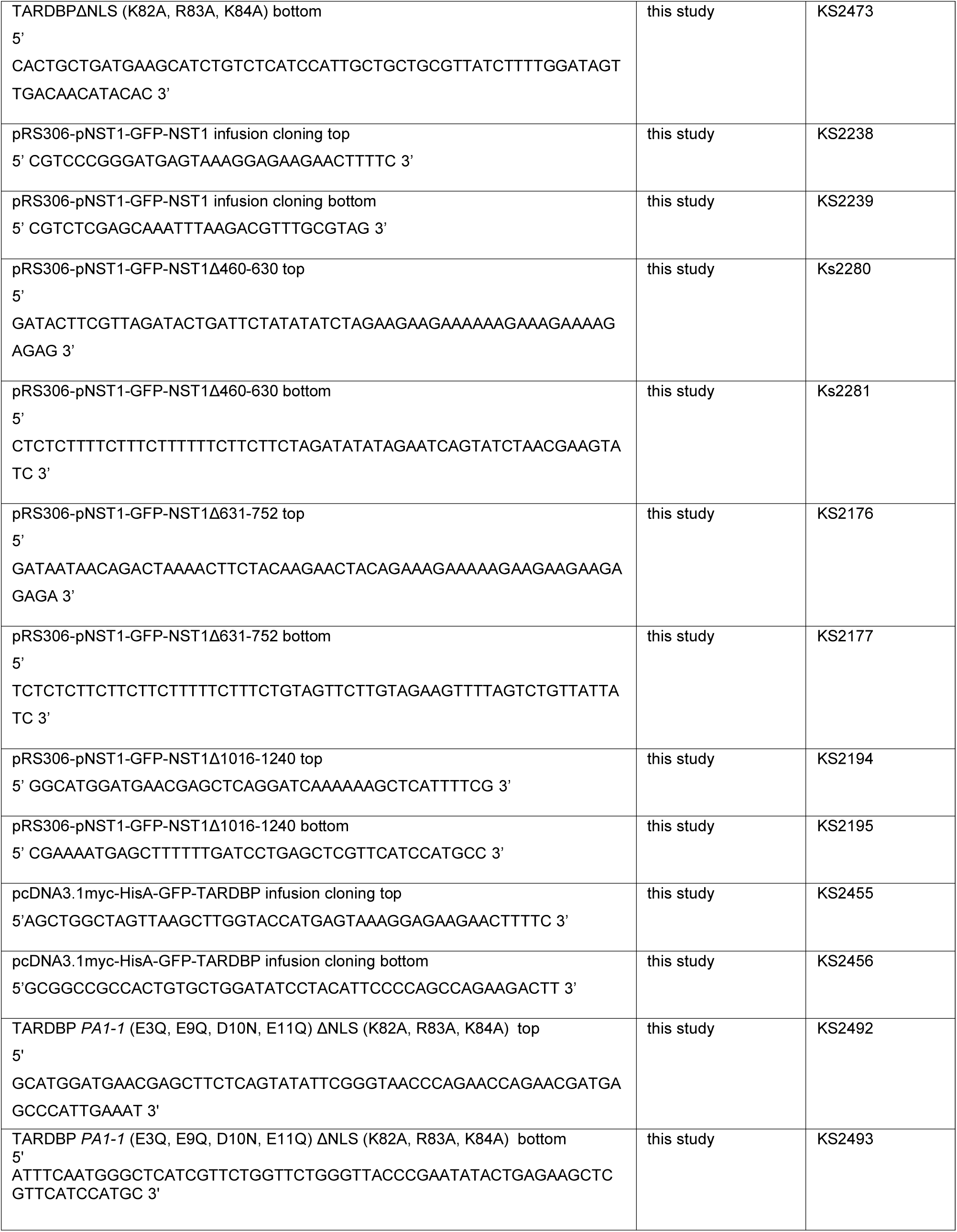
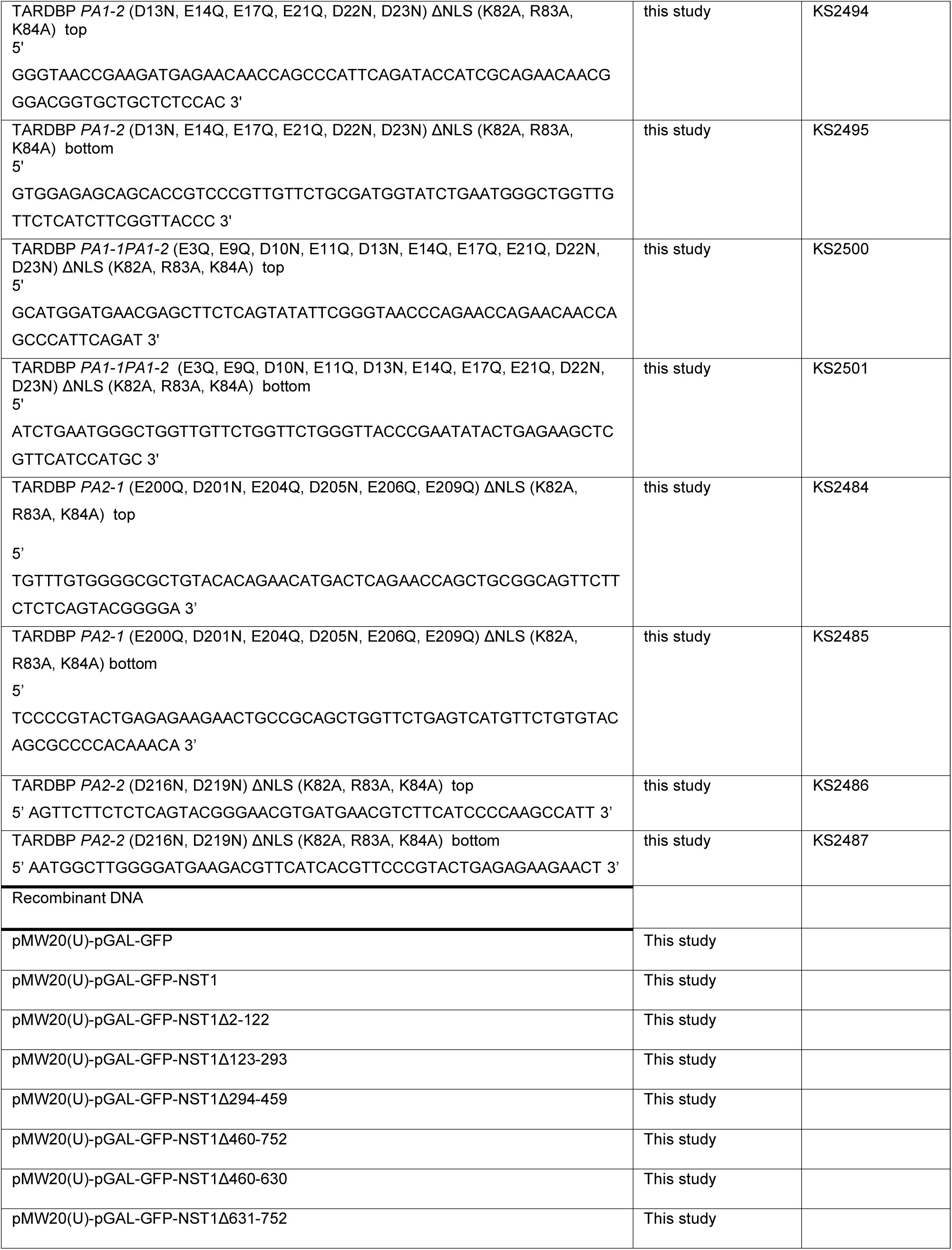
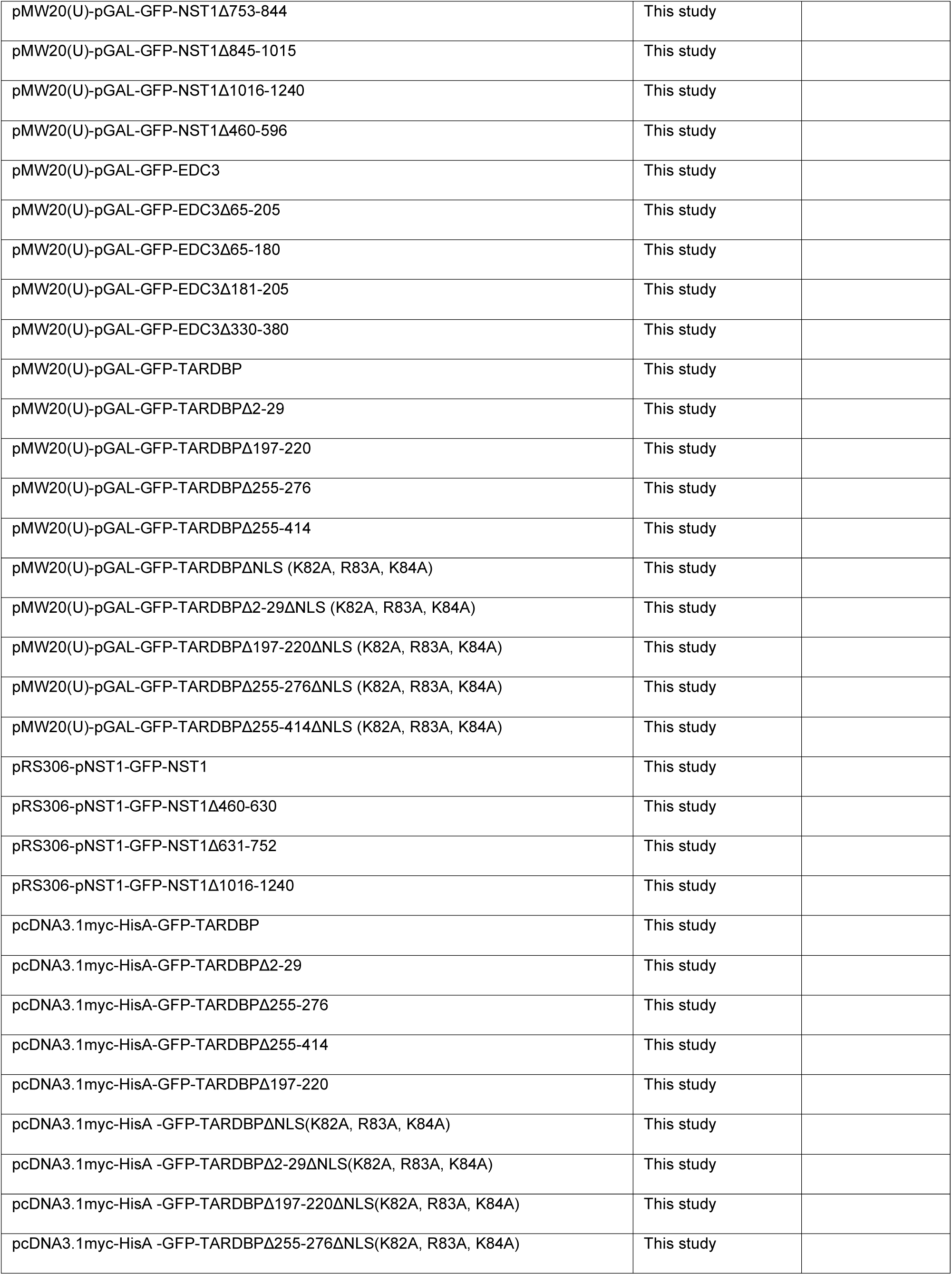
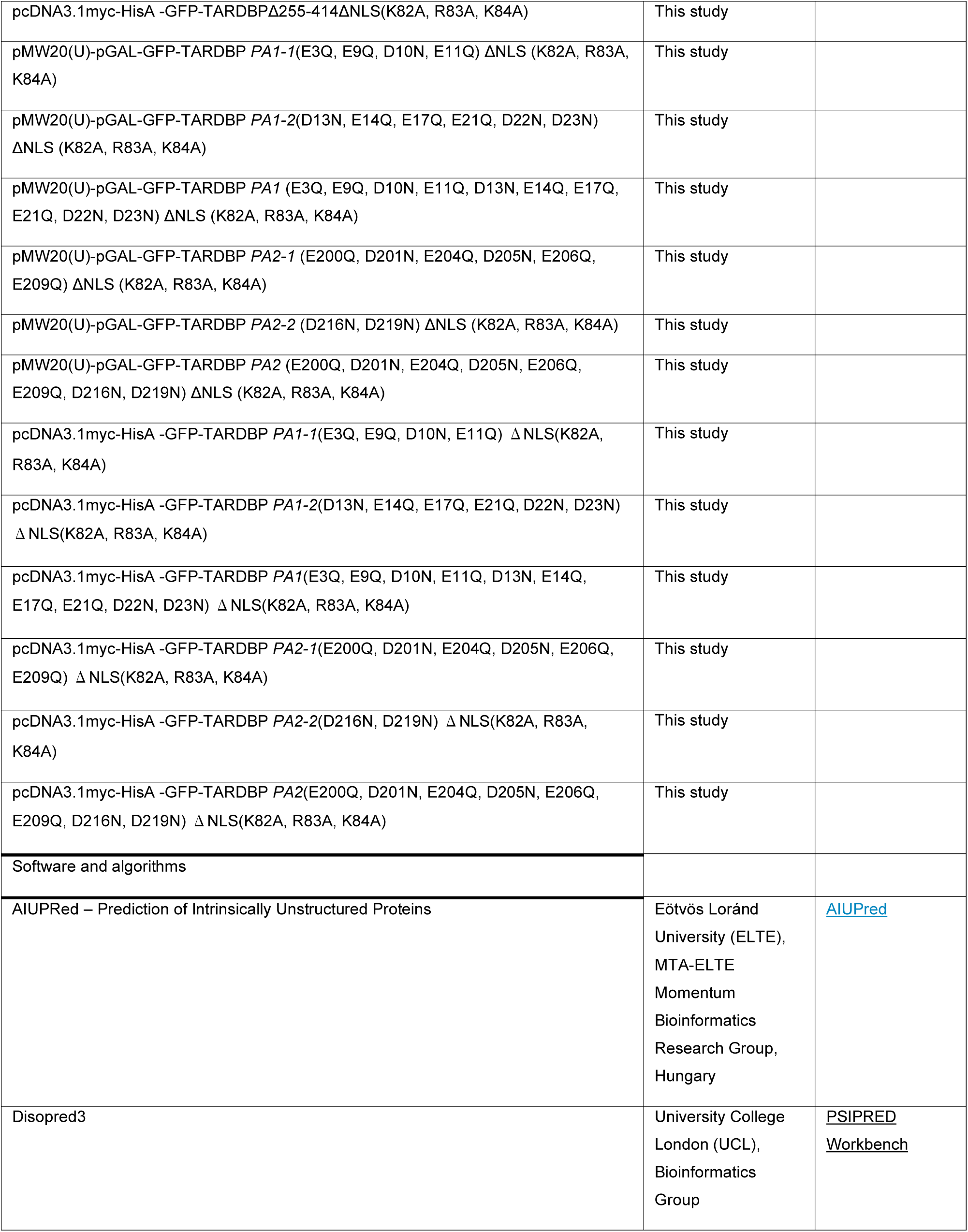
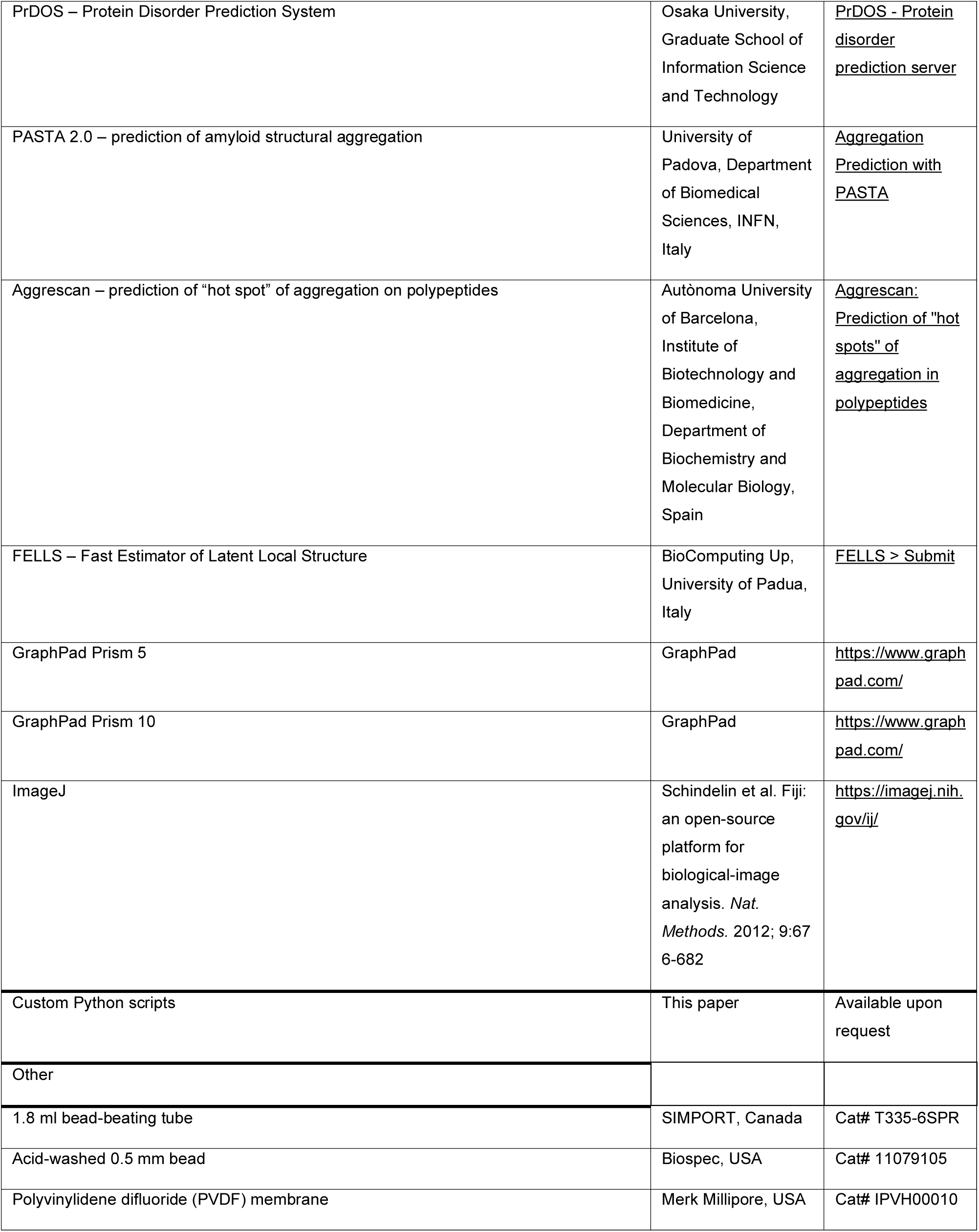

## METHOD DETAILS

### Yeast cell culture and transformation

*Saccharomyces cerevisiae* yeast cells were grown in YPAD medium until reaching the exponential growth phase (OD_600_ 0.5). Cells were harvested by centrifugation at 4,000 rpm for 5 min at room temperature (RT). The cell pellet was gently resuspended in 1 mL of 0.1 M LiOAc (Sigma-Aldrich, USA, L6883-250G) and incubated at RT for 5 min, followed by another centrifugation under the same condition. For transformation, cells were gently resuspended in 360 µL of yeast transformation mixture consisting of 50% polyethylene glycol (Sigma-Aldrich, USA, P4338-500G), 1 M LiOAc, 2 mg/mL salmon sperm DNA (Sigma, USA, D1626), and ddH_2_O. 100 ng plasmid DNA was added to the cell suspension, and the mixture was incubated at RT for 30 min. Heat shock was performed by incubating the cells at 42°C for 30 min, followed by cooling at RT for 3 min. Cells were harvested again by centrifugation at 4,000 rpm for 5 min at RT, and plated onto SC-Ura plates. Plates were incubated at 25°C until colonies appeared typically 2–3 days.

### Protein overexpression in yeast

The cells transformed with plasmids containing NST1, EDC3, *TARDBP* and their deletion mutants were cultured in a SC-U + 2% glucose medium up to the exponential growth phase (OD_600_ 0.5) and harvested. The cells were washed with a SC-U + 2% raffinose + 0.1% glucose medium three times, diluted to half its concentration, and cultured for an additional 3 hours in SC-U + 2% raffinose + 0.1% glucose. Thereafter, galactose was added to a final 2% concentration and incubated for designated hours.

### 1,6-Hexanediol treatment of yeast

Yeast cells induced for protein overexpression in 2% galactose for 4 hours were harvested by centrifugation at 4000 rpm for 5 minutes, and the resulting pellet was collected. The cell pellet was then resuspended and incubated in SC-U medium supplemented with 2% galactose, 2% raffinose, 0.1% glucose, and 10% 1,6-hexanediol (Daejung Chemical & Metals Co., LTD, South Korea, 629-11-8) for 1 hour.

### Chromosomal integration of genes in yeast

Various NST1 deletion mutants were acquired by PCR-mediated deletion cloning with the full length pRS306-pNST1-NST1 and verified by sequencing. Plasmids pRS306-pNST1-NST1 and its deletion mutants were treated with EcoRV to make a single cut at the URA3 and were transformed into *Δnst1* cells (YSK3490) by 42 °C heat shock. Integration into the chromosome occurred at low frequency via homologous recombination at the URA3. The transformed cells were plated on the SC plate without uracil and colony PCR was performed to confirm whether the fragment was successfully transformed.

### Infusion cloning of genes

The PCR-amplified coding sequences were generated using primers containing 15-bp extensions homologous to the vector ends. The destination vector pcDNA3.1-myc-His was linearized by restriction enzyme digestion at 37°C for 3 h. The digested vector and PCR fragments were then subjected to infusion cloning reaction (EZ-HT cloning kit, Enzynomics, Korea, EZ016TM), which utilizes single-strand annealing activity to join DNA fragments with complementary ends, circumventing the need for conventional ligation. The reaction products were transformed into chemically competent *E. coli* DH5α cells, and transformants were selected on LB agar plates supplemented with ampicillin (100 μg/ml). The integrity of the resulting constructs was verified by DNA sequencing.

### TDP-43 PA1 and PA2 Domain Mutagenesis

PCR-based site-directed mutagenesis was performed using the plasmid pMW20(U)-pGAL-GFP-TARDBPΔNLS (K82A, R83A, K84A) and pcDNA3.1-myc-hisA-GFP-TARDBPΔNLS (K82A, R83A, K84A) as a template to introduce targeted amino acid substitutions within the TDP-43 PA1 and PA2 domain. The *PA1-1* mutant was generated with primers KS2492 and KS2493, and the *PA1-2* mutant with KS2494 and KS2495. The *PA2-1* mutant was generated with primers KS2484 and KS2485, and the *PA2-2* mutant with primers KS2486 and KS2487. The *PA1-1PA1-2* and *PA2-1PA2-2* mutants, which harbor both the *PA1-1* and *PA1-2* mutations or *PA2-1* and *PA2-2* mutations, were constructed using the *PA1-2* or *PA2-1* plasmid as a template with the *PA1-1PA1-2* primers (KS2500 and KS2501) or the *PA2-2* primers (KS2486 and KS2487), respectively. The sequences of all primers were listed in the KEY RESOURCES TABLE. All mutant strains were verified by DNA sequencing.

### Widefield microscopy of cells and image analysis

GFP-labeled proteins expressed in the cells were visualized using an Axioplan2 microscope (Carl Zeiss,Jena, Germany) equipped with a 100x Plan-Neofluar oil immersion objective. Consistent culture conditions, exposure times, and fluorescence intensities were maintained across all strains in this study to ensure a comparative analysis of puncta intensity. Image analysis was performed using FIJI ImageJ (https://imagej.nih.gov/ij/).

### Confocal microscopy of cells and image analysis

GFP-labeled proteins expressed in the cells were imaged using both a ZEISS LSM900 laser scanning confocal microscope (Carl Zeiss, Jena, Germany) equipped with high-sensitivity GaAsP-PMT detectors (2 standard and 1 additional GaAsP-PMT), and a ZEISS LSM980 confocal microscope with an Airyscan 2 detector (Carl Zeiss, Jena, Germany). For Airyscan super-resolution imaging with the LSM980, the Airyscan Joint Deconvolution mode was utilized, achieving lateral resolution down to ∼90 nm. Both systems were fitted with a 63x/1.4 NA oil immersion objective, and all images were acquired at room temperature. Microscope settings, including laser power, detector gain, and pinhole size, were kept identical for all samples to ensure quantitative comparability. Image acquisition and processing were performed using ZEISS Zen software (version 3.9, Carl Zeiss).

### FCS measurements and data analysis

All FCS measurements were performed at 25°C on a confocal microscope (LSM780; Carl Zeiss, Inc. Germany) combined with a ConfoCor3 and a 40×/1.2 NA water-immersion objective (Carl Zeiss, Inc.). Details of the analysis of fluorescence auto-correlation functions (FAFs) were described in the previous study (Kawai-Noma et al., 2006; Pack et al., 2014), FAFs were fitted to a two-component model with a triplet term.

For each cell, the cytoplasmic region was identified by visual inspection of the fluorescence image, avoiding the GFP visualized puncta. At least five independent measurements (each lasting 10 s) were recorded per cell. Over 15 cells were analyzed per condition.

Autocorrelation curves were generated using ZEN black software (Carl Zeiss, Inc.) and fitted with two-component 3D diffusion model including a triplet term:

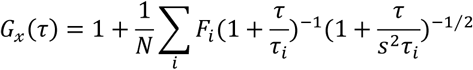

Where 𝐹_𝑖_ and 𝜏_𝑖_ are the fraction and diffusion time of component I, respectively; 𝑁 is the average number of fluorescent molecules in the confocal volume defined by the radius 𝑤_0_ and the length 2𝑧_0_; and 𝑠 is the structure parameter representing the ratio 𝑠 = 𝑧_0_⁄𝑤_0_ (axial/lateral radius ratio of the confocal volume). The structural parameter was calibrated using a known concentration of Rh6G standard solution at room temperature. The relationship between absolute concentration and calculated N was calibrated using a known concentration of Rh6G or recombinant mGFP solution. The effect of photobleaching on evaluation of concentration was determined according to a previous study. Fluoprescence correlation functions (FCFs) in live cells were measured sequentially 6 times with duration of 10s to minimize photobleaching effect.

The cytoplasmic concentration of the target protein was calculated as

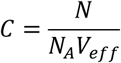

Where 𝑁_𝐴_ is Avogadro’s number and 𝑉_𝑒𝑓𝑓_ is the effective confocal volume determined from calibration measurements with Rhodamine 6G solutdion.

### Fluorescence recovery after photobleaching (FRAP) in living yeast cells

Fluorescence recovery after photobleaching (FRAP) was performed on puncta induced by GAL promoter using a Zeiss LSM900 laser scanning confocal microscope with a 100×/1.40 NA oil immersion objective (Carl Zeiss AG, Oberkochen, Germany). Cells were mounted on agarose pads, composed of SC 2% raffinose/ 2% galactose media and 1∼3% agarose, to minimize the delicate movements of cells. For the confocal imaging, 0.6% 488 nm wavelength laser and GaAsP-PMT detector were used. Due to the low GFP signals of proteins and the small size of yeast, the pinhole was opened to 3 Airy units, about 240 μm. For fast imaging, the acquisition mode was set to 512 px X 512 px frame size, 8 bits per pixel, 9 (about 1 μs of pixel times) scan speed, one-way, and none averaging scanning. For the FRAP experiments, the region of interest (ROI) was selected, and 200 cycles with no interval of time series were set. After snapping five images, bleaching on ROI was performed repeatedly three times with the power of 10∼100% 488 nm laser beam. Bleached signals on ROI were recovered, and these signals were acquired for the left 194 cycles. All the experiment was repeated about 20∼30 foci of each strain. If the foci were out of focus for 200 cycles, these data were discarded.

### Image analysis for FRAP

The mean intensity in ROI at each cycle was analyzed and drawn as graphs using Fiji with the Stowers plugin. All the graphs were normalized to 0-1, and the average curves of these graphs were calculated with GraphPad Prism 10 (GraphPad Software, CA, USA). These normalized curves were fitted with Eq.1.

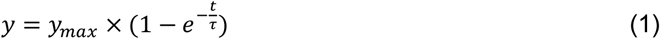

In Eq.1, 𝑦_𝑚𝑎𝑥_ is the 𝑦 value at infinite time (𝑡 ≈ ∞), and 𝜏 (tau) is the time point when the 𝑦 value is equal to 63.2% of 𝑦_𝑚𝑎𝑥_ value. The first derivative functions of the fitted curves were analyzed with GraphPad Prism 10 to calculate the maximum velocity of the recovery curves.

### Fixation and DAPI staining of yeast cells

Following 4 hours of galactose-mediated induction, yeast cells were harvested by centrifugation at 4,000 rpm for 5 minutes at RT. The supernatant was carefully removed, and cell pellets were resuspended in 4% paraformaldehyde (PFA) dissolved in Dulbecco’s phosphate-buffered saline (DPBS) (T&I, South Korea, BPP-9004) for fixation by gentle inversion mixing for 1 h at RT. To remove residual PFA, cells were subjected to two successive wash cycles consisting of centrifugation (4,000 rpm, 2 minutes) and resuspension in fresh DPBS (Welgene, South Korea, LB 001-02). The final cell pellet was reconstituted in DPBS and evenly distributed onto microscope slides. Vectashield mounting medium for fluorescence with DAPI (Vector Laboratories, Inc., USA, H-1200) was applied to each sample, followed by gentle mixing using a pipette tip to ensure homogeneous distribution of cells. Cellular morphology and fluorescence signals were subsequently analyzed using a Axioplan2 microscope (Carl Zeiss, Jena, Germany) equipped with a DAPI-specific filter set.

### U2OS cell culture and transfection

U2OS cells (ATCC, CRL-1427, HTB 96, Manassas, VA, USA) were cultured in Dulbecco’s Modified Eagle Medium (DMEM with high glucose; Cytiva, USA SH30023.01) supplemented with 10% fetal bovine serum (FBS; Cytiva, USA, SH30088.03) and 1X antibiotic-antimycotic solution (Anti-Anti; Gibco, USA,15240-062) at 37°C in a humidified incubator with 5% CO₂. Once the cells reached 70% confluence, they were transfected with pcDNA3.1-myc-hisA expression vectors encoding GFP-TDP-43, its IDR deletion mutants and charge neutralized mutants using Lipofectamine 3000 (Invitrogen, USA, L300015), following the manufacturer’s protocol. After transfection, the cells were selected with 600 ng/μL Geneticin™ Selective Antibiotic (G418 Sulfate) (Gibco, USA, 10131035).

### Fixation and DAPI staining of U2OS cells

Forty-eight hours after transfection, U2OS cells cultured on a cover slide were fixed with 4% paraformaldehyde (PFA, T&I, South Korea, BPP-9004) at RT for 1 h, washed three times with phosphate-buffered saline (PBS, WELGENE, South Korea, LB 001-02)., and mounted using a DAPI-containing mounting solution (Vector Laboratories, Inc., USA, H-1200). Fluorescence images were acquired using an Axioplan2 fluorescence microscope (Carl Zeiss, Jena, Germany) equipped with a 20x/0.5 Ph2 Plan-Neofluar objective.

### Western blotting analysis

Protein extraction from yeast cells was obtained using the alkaline lysis as described in Choi and Song (*Int. J. Mol. Sci. 2022*, *23*, 2501)U2OS cell extract was obtained as previously described (PLOS ONE | https://doi.org/10.1371/journal.pone.0302936 May 7, 2024). Protein samples were separated by sodium dodecyl sulfate–polyacrylamide gel electrophoresis (SDS-PAGE) and transferred onto polyvinylidene difluoride (PVDF) membranes (Merck Millipore, Burlington, MA, USA) using a semi-dry transfer system. Anti-GFP antibody (600-101-215 ROCKLAND, Limerick, PA, USA, 1:1000 dilution) was used for Western blotting. To confirm equal protein loading, the membranes were stripped of the anti-GFP antibody prior to re-probing with an anti-Tubulin antibody anti-Tubulin (Mouse, Sigma-Aldrich, Cat# T5168, 1:10000 dilution) and subsequently with HRP-conjugated anti-mouse secondary antibody (Santa Cruz Biotechnology, Cat# sc-2005, 1:10000 dilution).

## QUANTIFICATION AND STATISTICAL ANALYSIS

Only statistically analyzed data are shown as mean ± SEM. Statistical analyses were performed using GraphPad Prism. Statistical significance was determined using unpaired two-tailed Student’s t-test with Welch’s correction. P < 0.05 was considered statistically significant.

## DATA AND CODE AVAILABILITY

Custom algorithms were implemented in Python 3.10 by the authors. The analysis pipeline included modules written with NumPy and SciPy and was used to process experimental data shown in Figure 1. The code used in this study was written by the authors and is available from the corresponding author upon reasonable request.

## ADDITIONAL RESOURCES

N/A

## DISCLOSURE

A preprint version of this manuscript is available on bioRxiv (DOI:)

## ACKNOWLEDGMENTS

We thank the Cellular Imaging Core Facility at the ConveRgence mEDIcine research cenTer (CREDIT), Asan Medical Center, for technical support. The authors would like to thank Ewha Women’s University Fluorescence Core Imaging Center for valuable guidance in FRAP analysis. We specially appreciate the technical assistance provided by professor Dongmin Kang, the Director, and Research Technician Sinae Jang, of Ewha Fluorescence Core Imaging Center. TDP-43 gene clone was kindly provided by Dr. Hyung-Jun Kim in Korea Brain Research Institute. This work was in part supported by the National Research Foundation of Korea (NRF) grant provided to K. Song (RS-2023-00243284).

## AUTHOR CONTRIBUTIONS

Conceptualization, K.S. and Y.J.C.; methodology, K.S., Y.J.C, and C.P.; Investigation, Y.J.C, Y.L., S.C., and Y.P.; writing—original draft, K.S. and Y.J.C.; writing—review & editing, K.S., Y.J.C, C.P, and Y.J.; funding acquisition, K.S.; resources, K.S. and C.K.B.; supervision, K.S., and Y.J.C

## Supplementary Figures

**Supplementary Figure 1.**
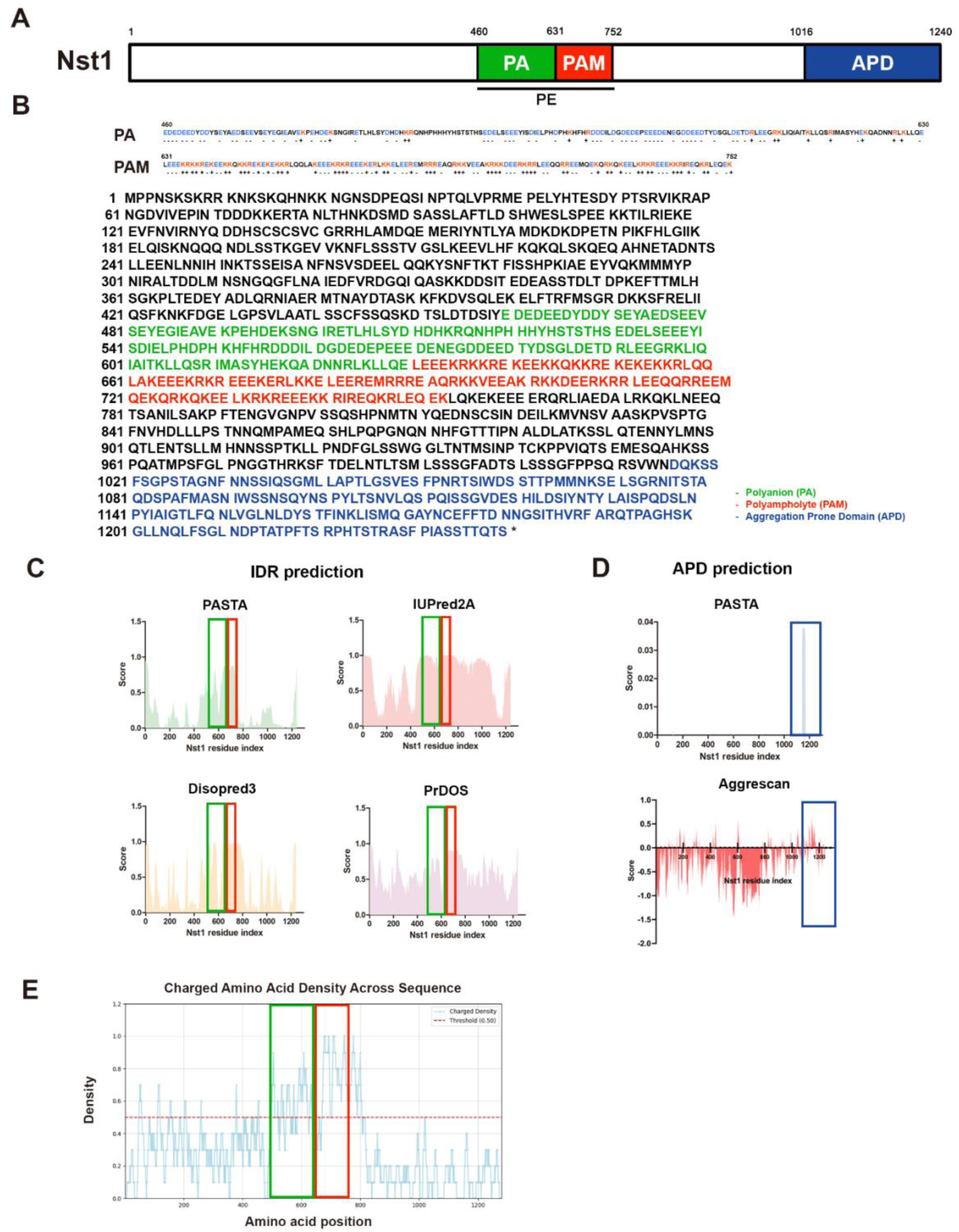
(A) A diagram of Nst1 showing domain architecture predicted by prediction tools. The PE region spanning residues 460–752 is indicated by a black line, while the PA region (residues 460–630), PAM region (residues 631–752), and APD region (residues 1016–1240) are denoted by green, red, and blue, respectively. (B) Diagram illustrating the full amino acid sequence of Nst1, highlighting the charge-rich regions designated as PA and PAM. In the PA and PAM sequence shown above, lysine (K) and arginine (R) residues, which are positively charged, are indicated in red with a (+) sign shown beneath each residue, while negatively charged glutamic acid (E) and aspartic acid (D) residues are indicated in blue with a (–) sign displayed below. In the full-length sequence, the PA, PAM, and APD regions are highlighted in green, red, and blue, respectively. (C-D) Data analysis and graph generation were conducted using GraphPad Prism 5. (C) Prediction of Intrinsically Disordered Regions (IDRs) using multiple computational tools: PASTA 2.0 (green), IUPred2A (pink), Disopred3 (yellow), and PrDOS (purple). Residues with scores > 0.5 consistently predicted by all four tools were classified as IDRs. The red box denotes the PAM region, while the green box indicates the PA region. (D) Prediction of Aggregation Prone Domains (APDs) utilizing PASTA 2.0 (blue) and Aggrescan (red). Regions exhibiting high positive scores were designated as APDs, as indicated by the blue box. (E) The density of charged amino acids along the Nst1 sequence was analyzed using a sliding window approach. A window of 10 consecutive amino acids was moved along each sequence in single-residue steps. For each window, the proportion of charged residues—defined as lysine (K), arginine (R), histidine (H), aspartic acid (D), and glutamic acid (E)—was calculated by dividing the number of these residues by the window size. Regions with a ratio equal to or greater than 0.5 were designated as high-density charged segments. The red and green boxes indicate the PAM and PA regions, respectively.

**Supplementary Figure 2.**
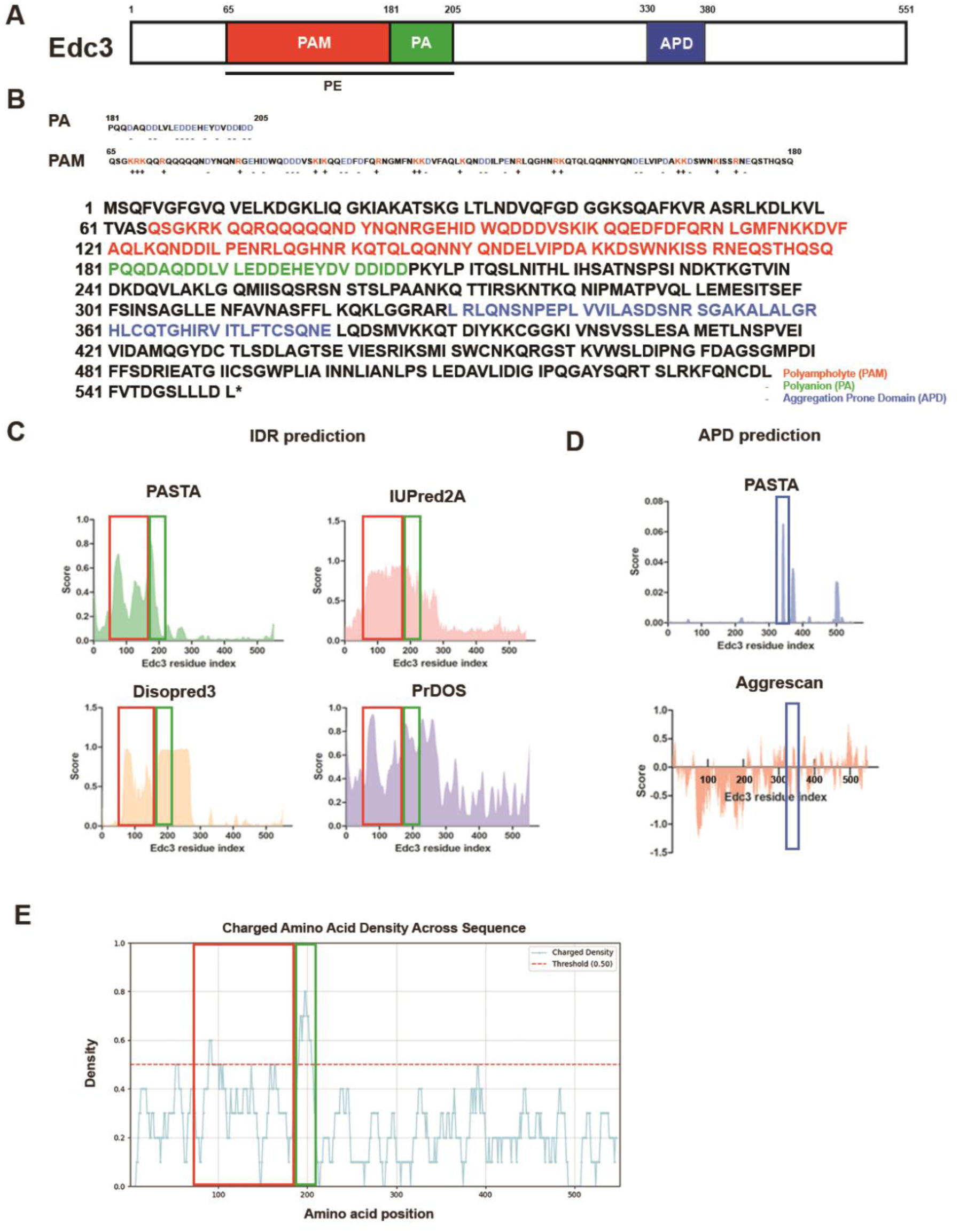
(A) Schematic representation of Edc3 domain organization as predicted by computational tools. The PE region (residues 65-205) is represented by a black line, whereas the PA (181-205), PAM (65-180), and APD (330-380) regions are highlighted in green, red, and blue, respectively. (B) Amino acid sequence map of Edc3, emphasizing regions rich in charged residues: the PA and PAM. The annotation of charged amino acids and the color scheme for the sequences were applied as in Supplementary Figure 1B. In the full-length amino acid sequence, the PA region is indicated in green, the PAM region in red, and the APD region in blue. (C-D) Graphical analysis and plotting were performed using GraphPad Prism 5. (C) Computational prediction of Intrinsically Disordered Regions (IDRs) in Edc3 using a suite of bioinformatics tools: PASTA 2.0 (green), IUPred2A (pink), Disopred3 (yellow), and PrDOS (purple). IDRs were defined as regions where all four algorithms consistently predicted disorder scores exceeding 0.5. The PAM and PA regions are demarcated by red and green boxes, respectively. (D) Identification of Aggregation Prone Domains (APDs) in Edc3 through analysis with PASTA 2.0 (blue) and Aggrescan (red). Regions displaying elevated positive scores were classified as APDs, highlighted by blue boxes. (E) The density of charged amino acids along the Edc3 sequence was analyzed using a sliding window approach. The detailed method is the same as described in Supplementary Figure 1E. The red and green boxes indicate the PAM and PA regions, respectively.

**Supplementary Figure 3.**
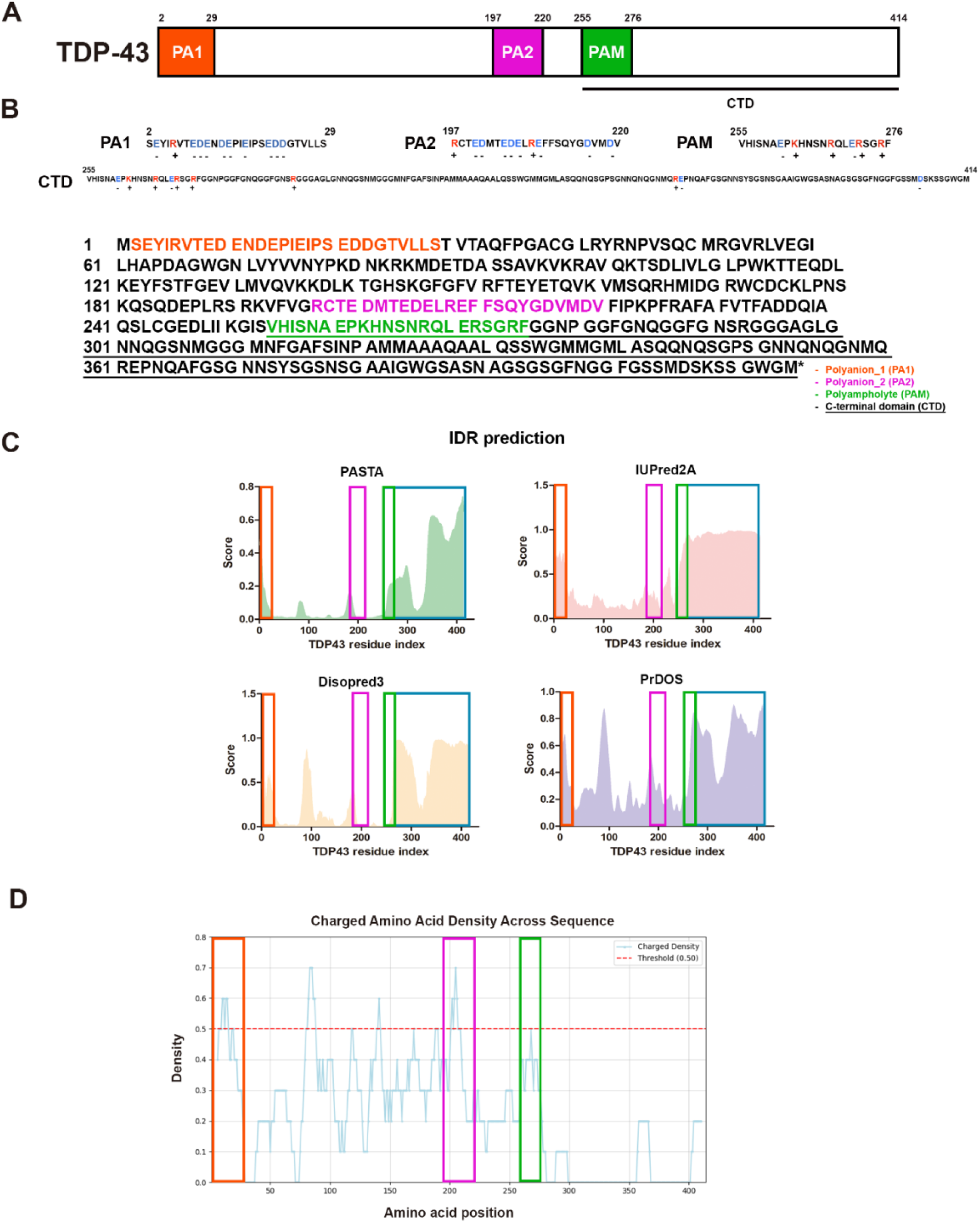
(A) A scheme of domain prediction of TDP-43. The CTD region, comprising residues 255–414, is depicted as a black line. The PA1 (2-29), PA2 (197-220), and PAM (255-276) regions are shown in red, pink, and green, respectively. (B) Amino acid sequence of TDP-43, highlighting the charged amino acid clusters (PA1, PA2 and PAM) and CTD. The annotation of charged amino acids was implemented in accordance with Supplementary Figure 1B. Each domain was indicated using the same box colors as in A, and the CTD is denoted with an underline. (C) Graphical representations were created and analyzed with GraphPad Prism 5. Computational analysis of Intrinsically Disordered Regions (IDRs) within TDP-43 employing multiple bioinformatics algorithms: PASTA 2.0 (green), IUPred2A (pink), Disopred3 (yellow), and PrDOS (purple). IDRs were defined as regions with disorder propensity scores above 0.2 and a high proportion of charged amino acids. The PA1 region is indicated by red boxes, and the PA2 region is indicated by pink boxes. The PAM region is indicated by green box, and the CTD is indicated by blue box. (D) The density of charged amino acids along the TDP-43 sequence was analyzed using a sliding window approach. The detailed method is the same as described in Supplementary Figure 1E. The PA1 region and the PA2 region is indicated by red box and pink box, respectively, while the PAM region is indicated by green box.

**Supplementary Figure 4.**
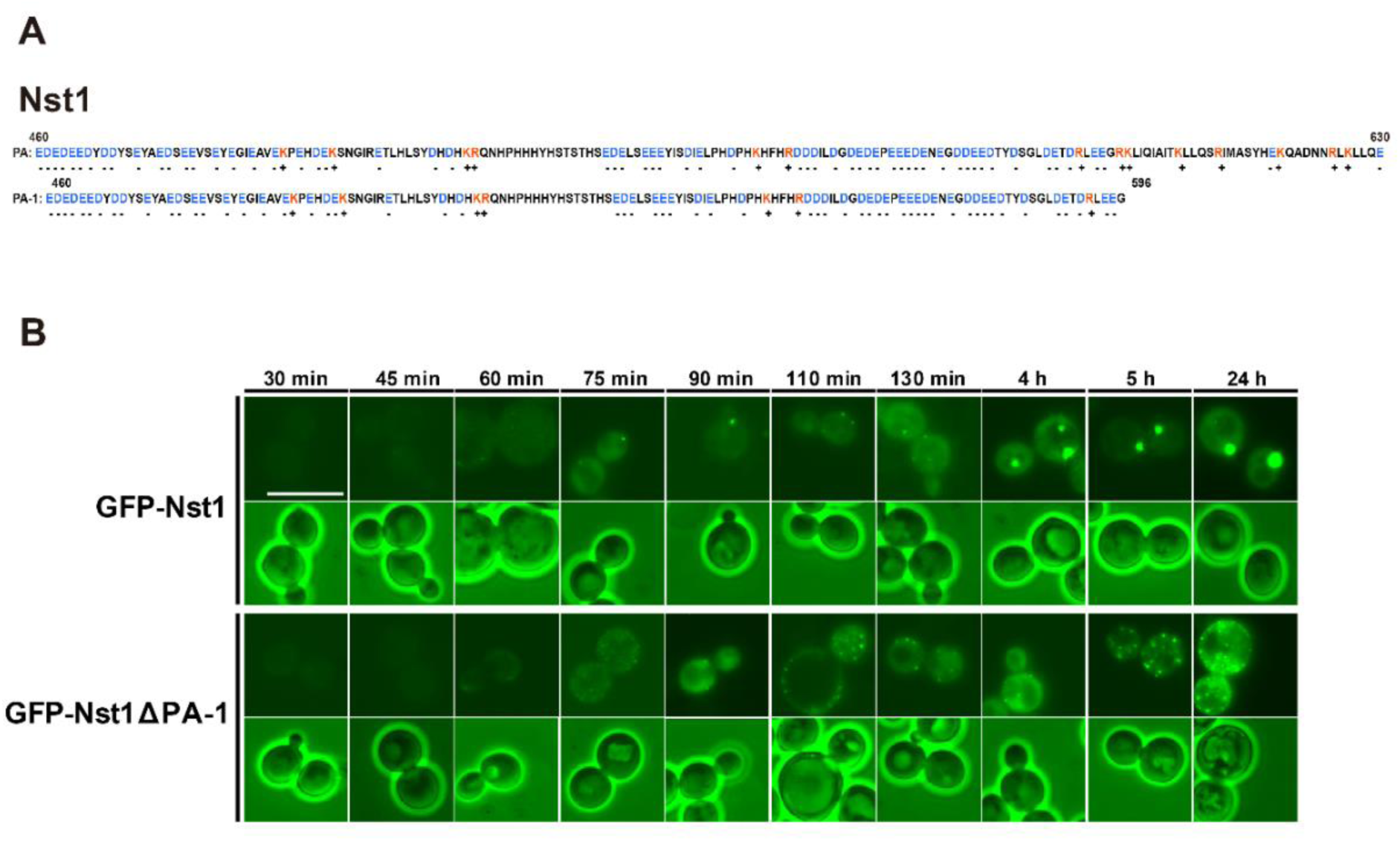
(A) Sequence of PA and PA-1 domains in Nst1 (B) Fluorescence microscopy images showing the formation of condensates by GFP-tagged Nst1 and Nst1ΔPA-1 in *Δnst1* (YSK3490) cells after GAL-induction. Images were acquired at the indicated time point following galactose induction. Scale bar: 10 μm.

**Supplementary Figure 5.**
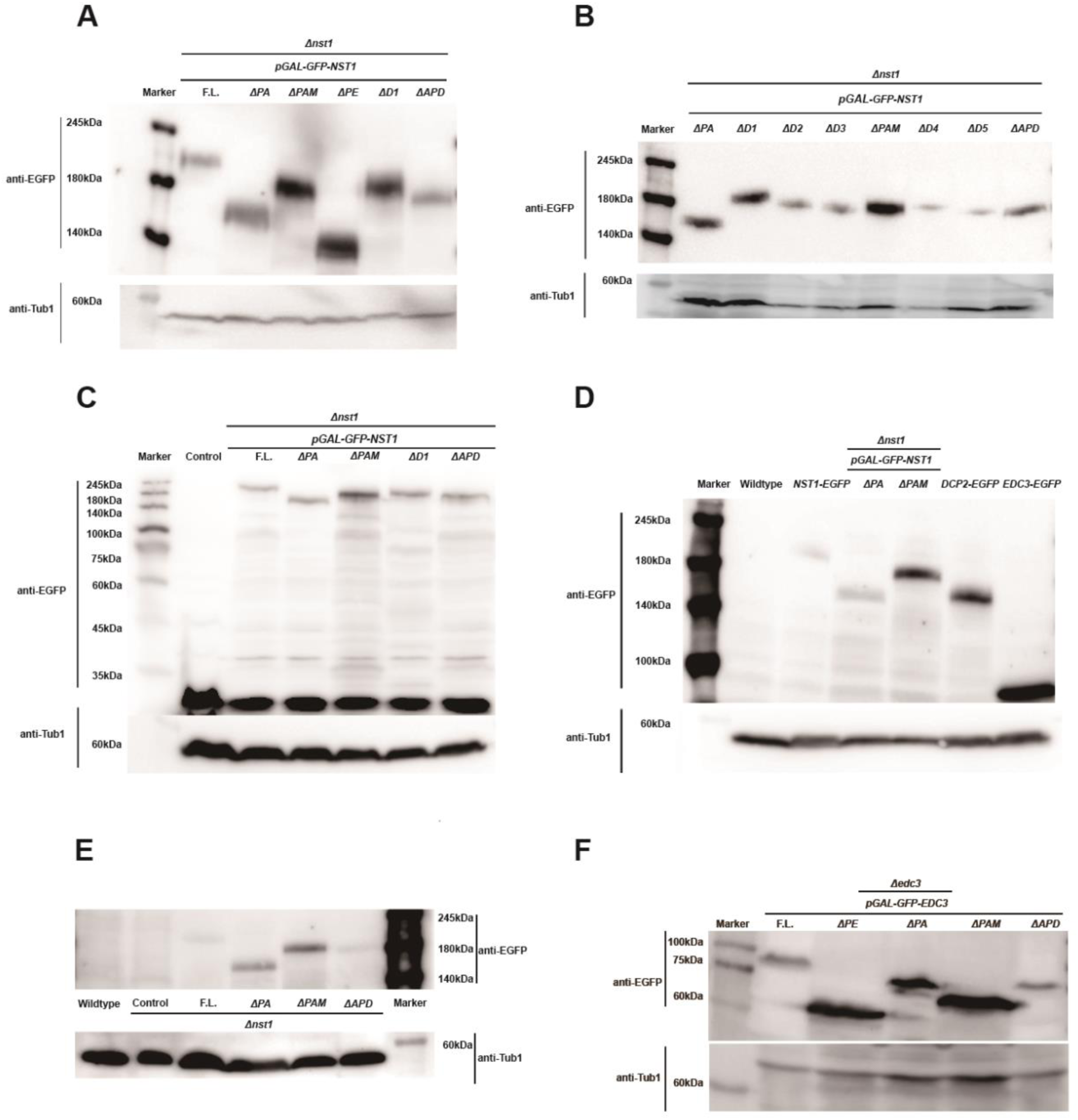
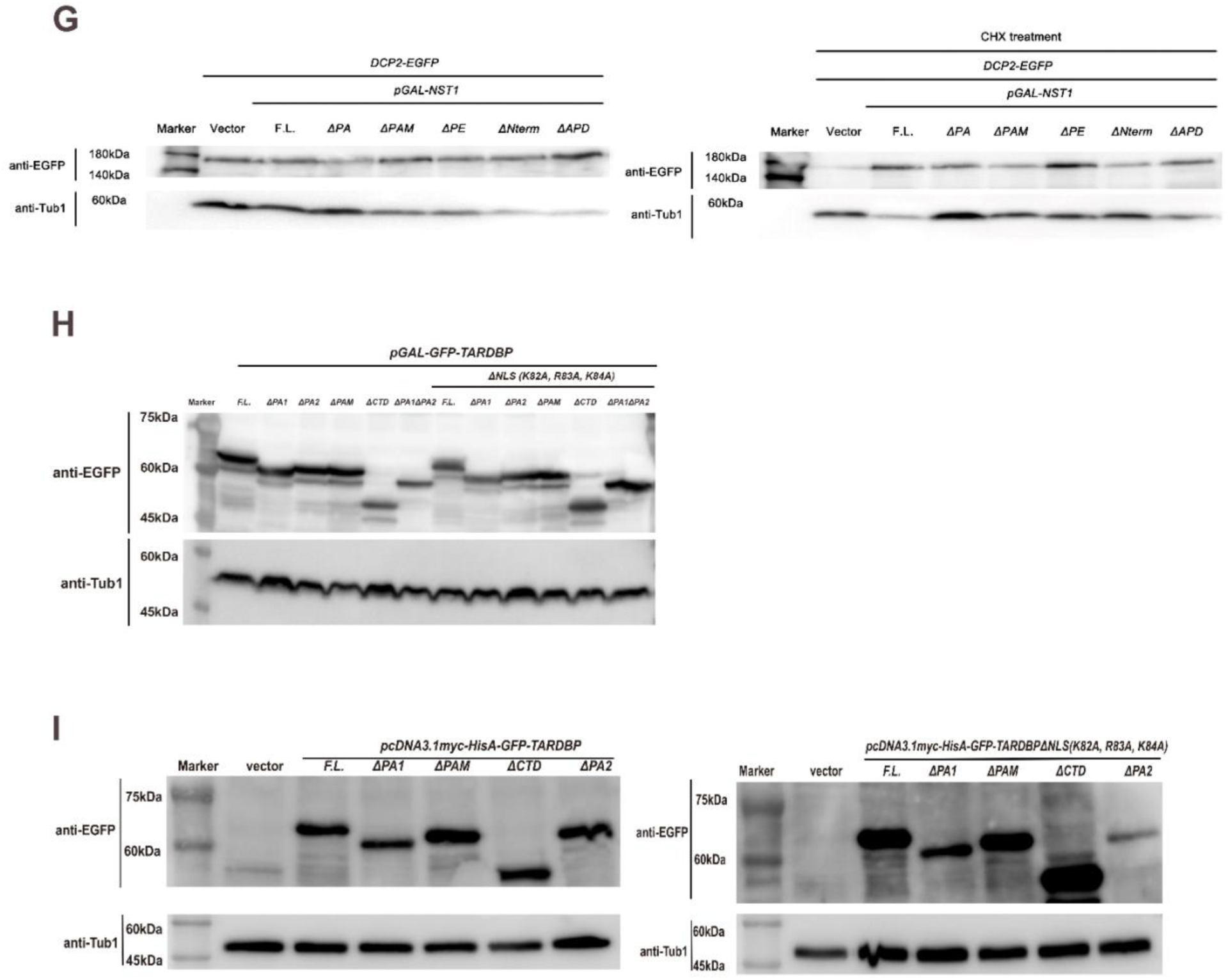
(A–B) Western blot analysis confirming the overexpression for GFP-tagged Nst1 domain deletion mutants, as shown in Figures 2C and 3B. Anti-GFP (600-101-215, Rockland) and anti-Tub1 (T5168, Sigma) antibodies were used to detect GFP-tagged fusion proteins and the Tub1 housekeeping protein, respectively. (A) The predicted molecular weights for Nst1 full-length, Nst1ΔPA, Nst1ΔPAM, Nst1ΔPE, Nst1ΔD1, Nst1ΔAPD, and Tub1 are 168.3 kDa, 148.1 kDa, 152.3 kDa, 132.1 kDa, 154.4 kDa, 144.1 kDa, and 49.8 kDa, respectively. (B) The predicted molecular weights for Nst1 full-length, Nst1ΔPA, Nst1ΔD1, Nst1ΔD2, Nst1ΔD3, Nst1ΔPAM, Nst1ΔD4, Nst1ΔD5, Nst1ΔAPD, and Tub1 are 148.1 kDa, 154.4 kDa, 148.7 kDa, 149.6 kDa, 152.3 kDa, 158.1 kDa, 149.9 kDa, 144.1 kDa, and 49.8 kDa, respectively. (C) Western blot verification of the overexpression of GFP-tagged Nst1 domain deletion mutants for FRAP analysis, as shown in Figure 4A. The predicted molecular weights for GFP (vector control), Nst1 full-length, Nst1ΔPA, Nst1ΔPAM, Nst1ΔD1, Nst1ΔAPD, and Tub1 are 26.8 kDa, 168.3 kDa, 148.1 kDa, 152.3 kDa, 154.4 kDa, 144.1 kDa, and 49.8 kDa, respectively. (D) Western blot analysis verifying protein expression prior to RNase inhibitor treatment, as shown in Figures 5G and 5I. The predicted molecular weights for Nst1-EGFP, Nst1ΔPA, Nst1ΔPAM, Dcp2-EGFP, Edc3-EGFP, and Tub1 are 168.3 kDa, 148.1 kDa, 152.3 kDa, 137.2 kDa, 88.1 kDa, and 49.8 kDa, respectively. (E) Western blot analysis confirming endogenous expression of GFP-tagged Nst1 full-length and deletion mutants (ΔPA, ΔPAM, ΔAPD) in log-phase *Δnst1* (YSK3490) cells, as shown in Figure 2H. Tub1 served as a loading control. The predicted molecular weights are: Nst1 full-length, 168.3 kDa; Nst1ΔPA, 148.1 kDa; Nst1ΔPAM, 152.3 kDa; Nst1ΔAPD, 144.1 kDa; Tub1, 49.8 kDa. (F) Western blot confirmation of the overexpression of GFP-tagged Edc3 domain deletion mutants, as shown in Figure 6B. The predicted molecular weights for Edc3 full-length, Edc3ΔPE, Edc3ΔPA, Edc3ΔPAM, Edc3ΔAPD, and Tub1 are 88.1 kDa, 71.2 kDa, 85.2 kDa, 74.2 kDa, 82.6 kDa, and 49.8 kDa, respectively. (G) Western blot analysis of Dcp2-EGFP (135.5 kDa) expression and Tub1 (49.8 kDa, loading control) in yeast cells used in Figure 5E. Cells were either untreated or treated with cycloheximide (CHX). (H) Western blot confirmation of the overexpression of GFP-tagged TDP-43 domain deletion mutants in budding yeast, as shown in Figure 6D. Constructs with intact nuclear localization signal (NLS) and corresponding NLS-deletion variants were analyzed. The predicted molecular weights are: TDP-43 full-length, 71.3 kDa; TDP-43ΔPA1, 68.1 kDa; TDP-43ΔPA2, 68.4 kDa; TDP-43ΔPAM, 68.7 kDa; TDP-43ΔCTD, 55.4 kDa. For NLS-deleted constructs: TDP-43ΔNLS full-length, 71.1 kDa; TDP-43ΔNLSΔPA1, 67.9 kDa; TDP-43ΔNLSΔPA2, 68.2 kDa; TDP-43ΔNLSΔPAM, 68.5 kDa; TDP-43ΔNLSΔCTD, 55.2 kDa. Tub1 served as a loading control (49.8 kDa). (I) Western blot analysis of pcDNAmyc-hisA-GFP*-TARDBP* constructs expressed in U2OS cells, as shown in Figure 6E. Both NLS-intact and NLS-deleted variants were examined. The predicted molecular weights are: TDP-43 full-length, 71.3 kDa; TDP-43ΔPA1, 68.1 kDa; TDP-43ΔPA2, 68.4 kDa; TDP-43ΔPAM, 68.7 kDa; TDP-43ΔCTD, 55.4 kDa. For NLS-deleted constructs: TDP-43ΔNLS full-length, 71.1 kDa; TDP-43ΔNLSΔPA1, 67.9 kDa; TDP-43ΔNLSΔPA2, 68.2 kDa; TDP-43ΔNLSΔPAM, 68.5 kDa; TDP-43ΔNLSΔCTD, 55.2 kDa. Tub1 served as a loading control (49.8 kDa).

**Supplementary Figure 6.**
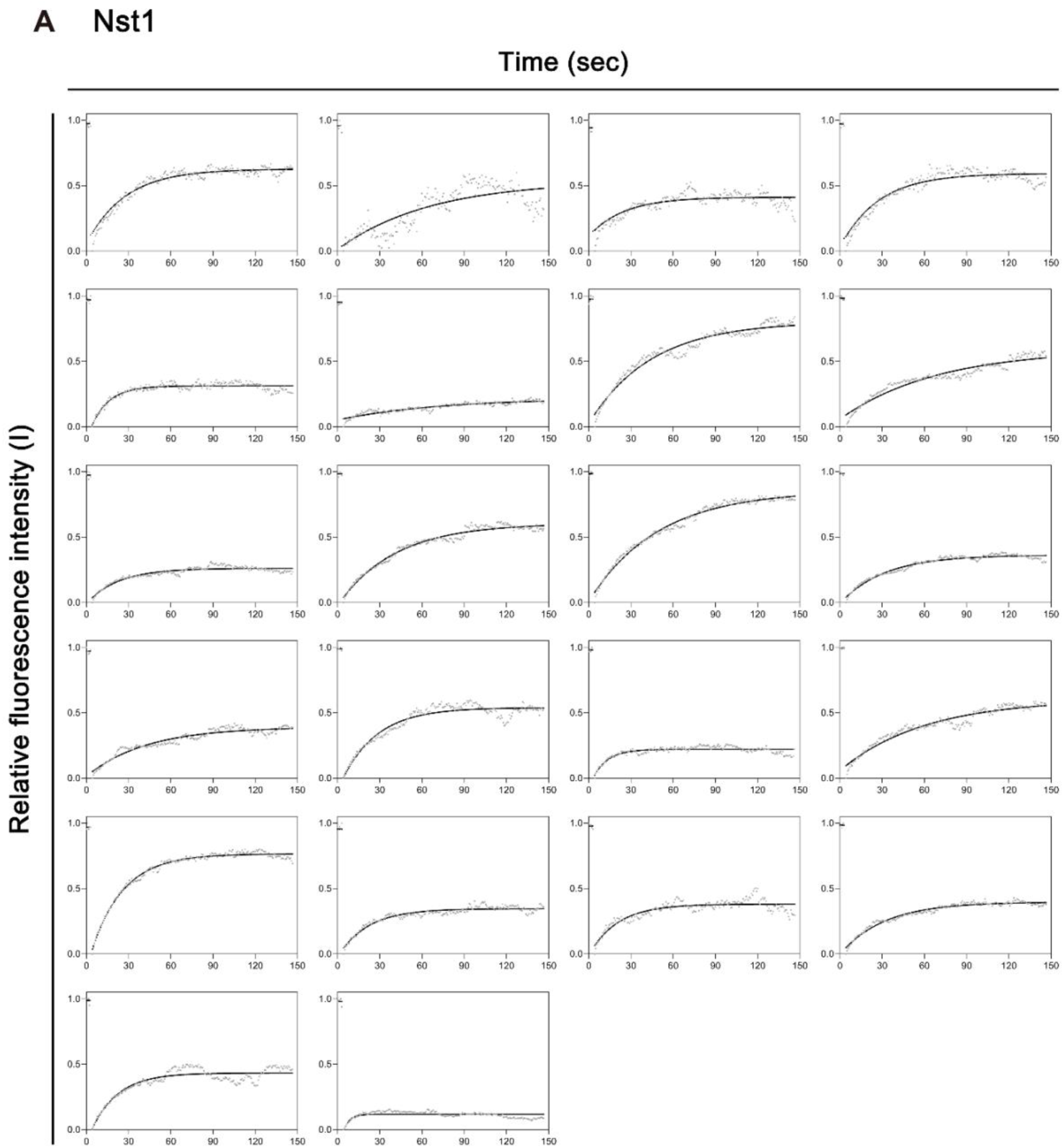
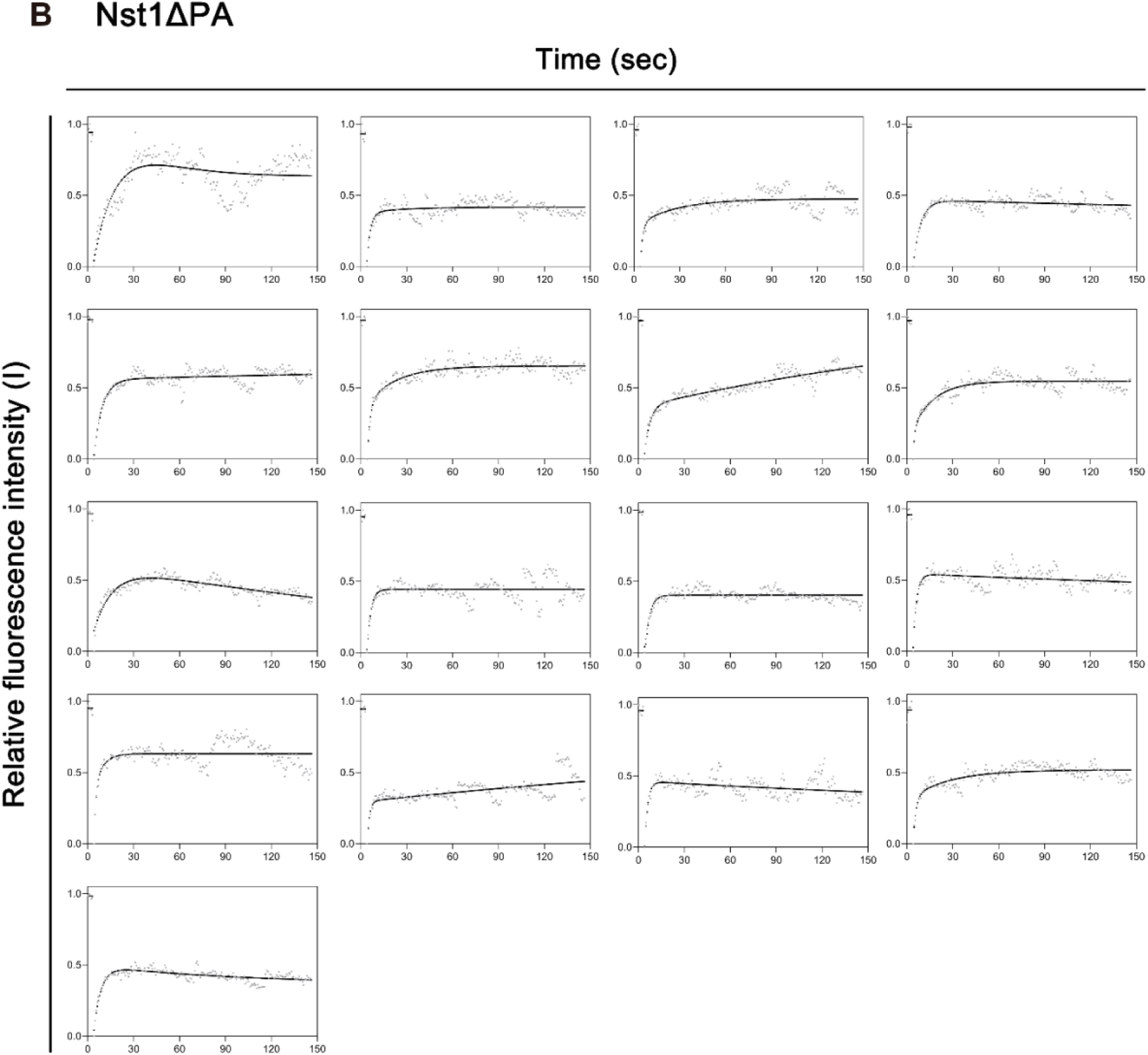
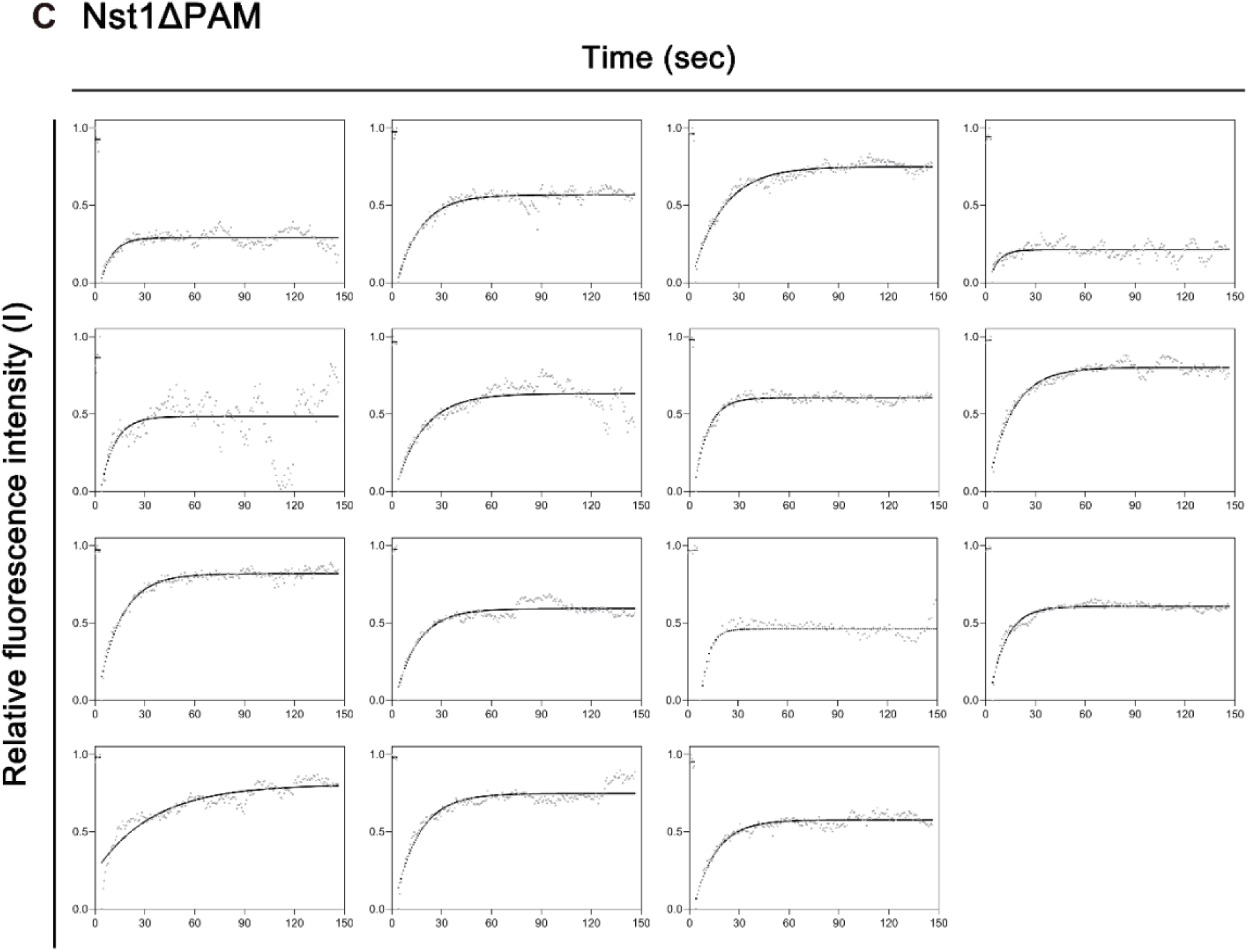
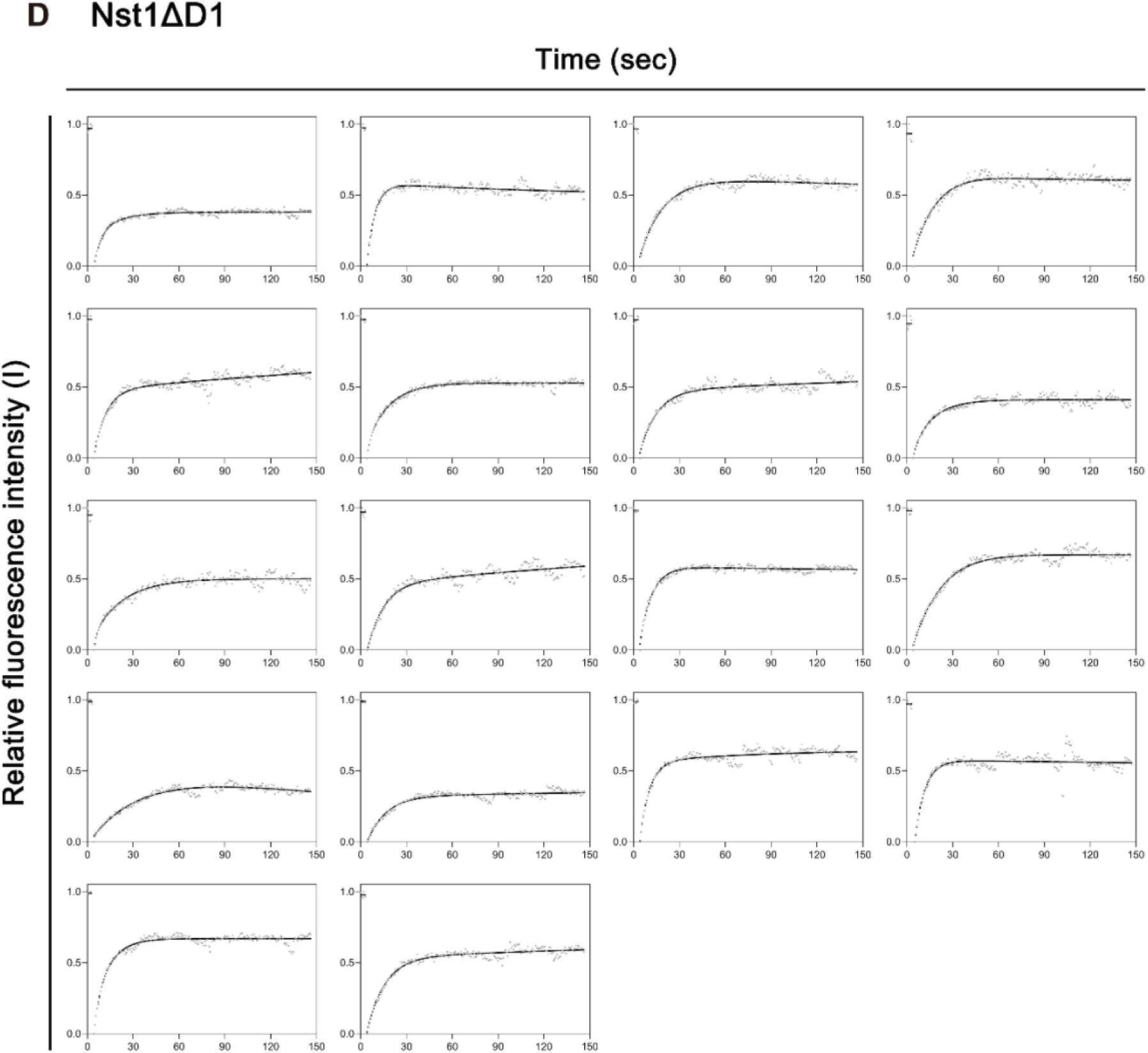
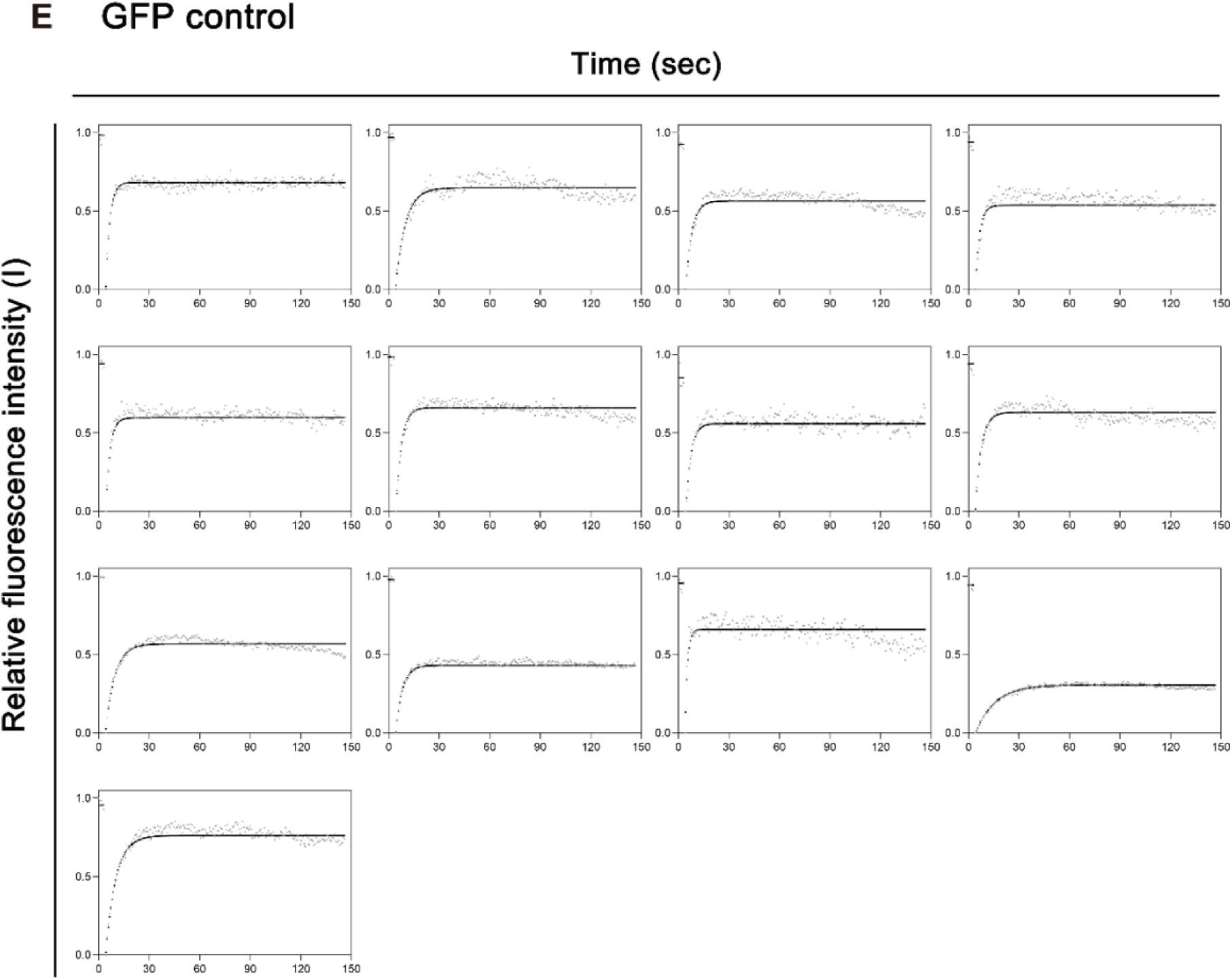

**Supplementary Figure 7.**
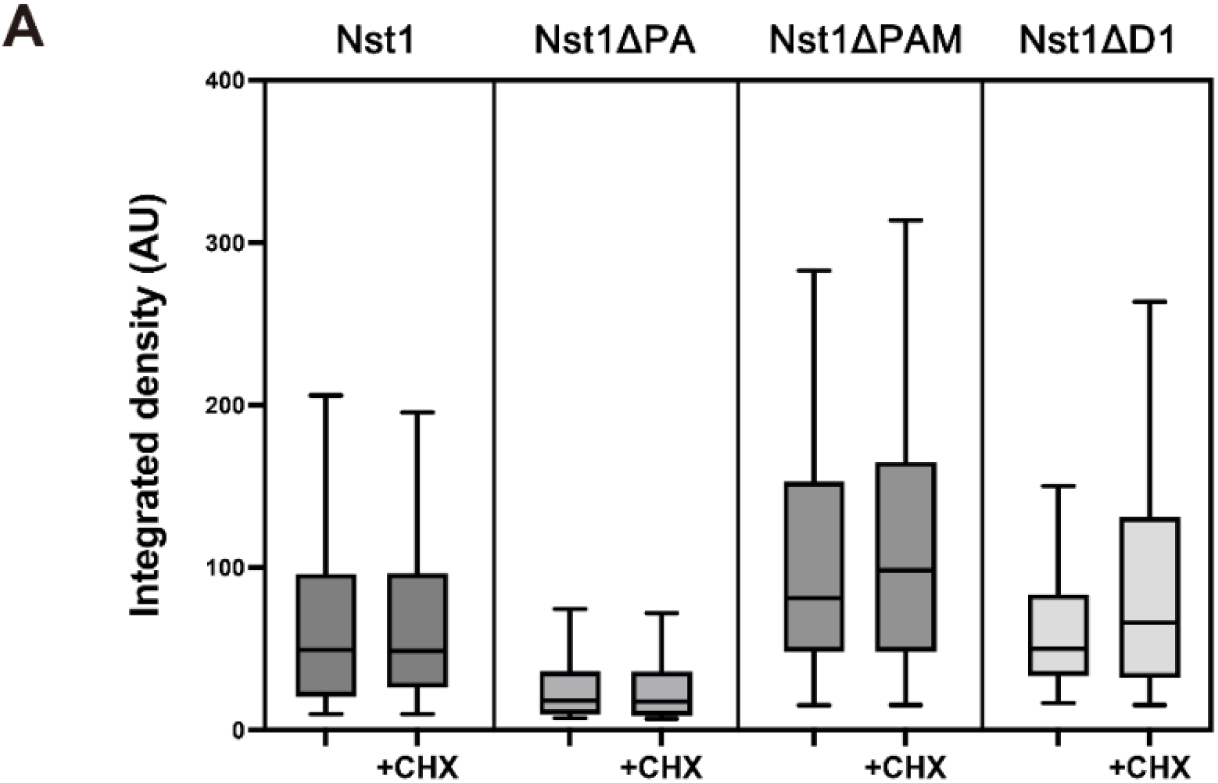
(A) Box plot of condensate formation, measured by integrated fluorescence intensity (Integrated Density), for condensates generated by galactose-induced, GFP-tagged Nst1, Nst1ΔPA, Nst1ΔPAM, and Nst1ΔD1 before and after CHX treatment. Fluorescence microscopy images were analyzed using Fiji (ImageJ) to quantify the integrated density of condensates. Data from more than 100 cells per condition were visualized using GraphPad Prism 10.

**Supplementary Figure 8.**
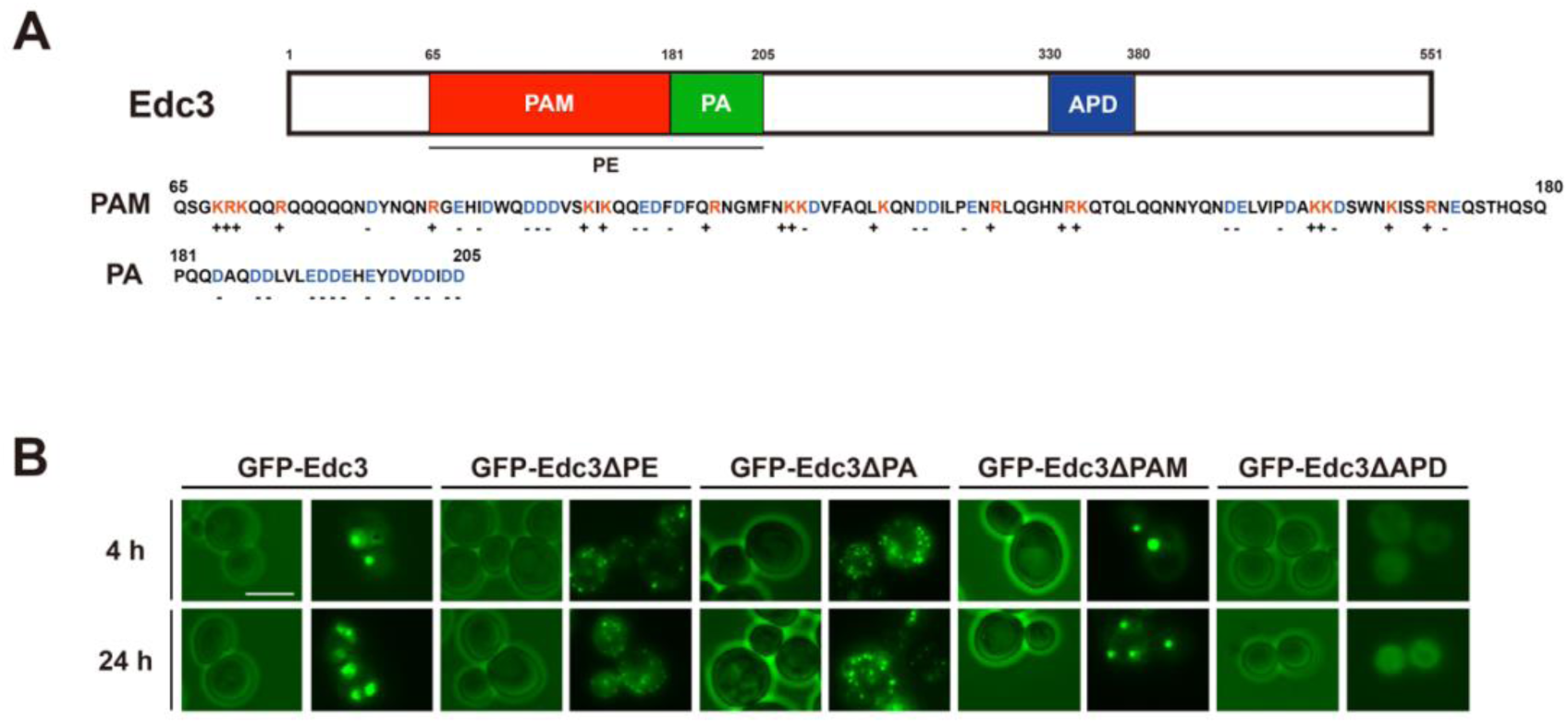
(A) Schematic representation of Edc3 domain architecture. The region spanning residues 65–205 was predicted to be the PE domain using a sliding window approach and various IDR prediction tools. See also Supplementary figure 2. The region from 330–380 was identified as the APD domain based on PASTA and Aggrescan analyses. The PE domain of Edc3 was further characterized according to its charge distribution: residues 181–205 exhibited a high density of negative charges and were thus designated as the PA domain. The remaining portion of the PE segment, encompassing residues 65–180 and containing a relatively balanced proportion of positively and negatively charged amino acids, was defined as the PAM domain. The charged amino acids in the charge-rich PAM and PA sequences were annotated as described in Figure 1D. (B) Fluorescence microscopy images showing condensate formation upon overexpression of Edc3, Edc3ΔPE, Edc3ΔPA, Edc3ΔPAM, and Edc3ΔAPD in *Δedc3* cells (YSK3707). Condensate formation for each Edc3 variant was imaged after 4 and 24 hours of galactose induction. For each strain and time point, representative images were selected from examinations of at least 100 cells.

**Supplementary Figure 9.**
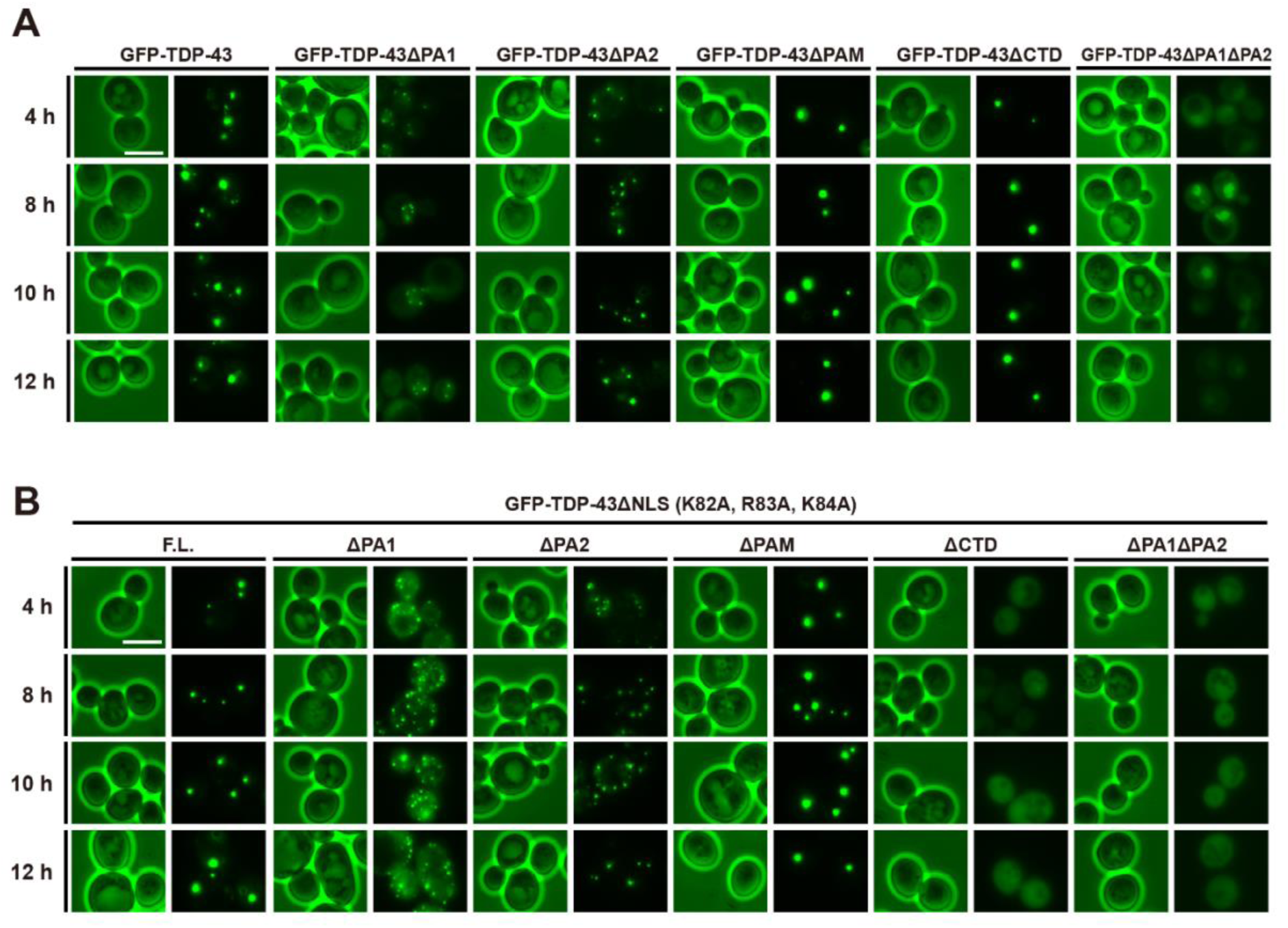
(A-B) Fluorescence microscopy images of budding yeast (W303a) cells after 4, 8, 10, 12 hours of overexpression of TDP-43 and its domain deletion mutants (ΔPA1, ΔPA2, ΔPAM, ΔCTD and ΔPA1ΔPA2) with or without NLS (Nuclear localization signal) sequence under a galactose-inducible promoter. Scale bar: 5 µm.

